# Co-habiting ants and silverfish display a converging feeding ecology

**DOI:** 10.1101/2023.11.17.567524

**Authors:** Thomas Parmentier, Rafael Molero-Baltanás, Catalina Valdivia, Miquel Gaju-Ricart, Pascal Boeckx, Piotr Łukasik, Nicky Wybouw

**Author notes:** **Correspondence** Thomas Parmentier, Nicky Wybouw.

## Abstract

1. Species from a great number of animal taxa have specialized to living with social hosts. Depending on their level of specialization, these symbiotic animals are characterized by distinct behavioural, chemical, and morphological traits that enable close heterospecific interactions. Despite its functional importance, our understanding of the feeding ecology of specialized animals living with social hosts remains limited.
2. In this study, we examined how host specialization of silverfish co-habiting with ants affects several components of their feeding ecology. We combined stable isotope profiling, feeding assays, phylogenetic reconstruction, and microbial community characterization of the *Neoasterolepisma* silverfish genus and a wider nicoletiid and lepismatid silverfish panel where divergent myrmecophilous lifestyles are observed.
3. Stable isotope profiling (δ^13^C and δ^15^N) showed that the isotopic niches (proxy for trophic niches) of granivorous *Messor* ants and *Messor-*specialized *Neoasterolepisma* exhibit a remarkable overlap within an ant nest. Trophic experiments and gut dissections further supported that these specialized *Neoasterolepisma* silverfish transitioned to a diet that includes plant seeds. In contrast, the isotopic niches of generalist *Neoasterolepisma* silverfish and generalist nicoletiid silverfish were clearly different from their ant hosts within the shared nest environment.
4. The impact of the myrmecophilous lifestyle on feeding ecology was also evident in the internal silverfish microbiome. Compared to generalists, *Messor-*specialists exhibited a higher bacterial density and a higher proportion of heterofermentative lactic acid bacteria. Moreover, the nest environment explained the infection profile (or the 16S rRNA genotypes) of *Weissella* bacteria in *Messor*-specialized silverfish and the ant hosts.
5. Together, we show that social hosts are important determinants for the feeding ecology of symbiotic animals, and can induce diet convergence.

## Introduction

Symbiosis, a close association between distinct organisms, is a pervasive ecological interaction (Paracer & Ahmadjian, 2000). Symbionts greatly vary in their strategies to interact with their host. These strategies have significant implications not only for the evolutionary ecology of the symbionts themselves but also for their hosts (Bronstein, 2015; Futuyma & Moreno, 1988). An initial differentiation among symbiont strategies can be made by considering their level of host dependency. Facultative species can survive independently but may benefit from the association with a host, whereas obligate symbionts are completely dependent on the symbiotic associations for survival and reproduction (Luong & Mathot, 2019; Poulin, 2011; Stadler & Dixon, 2005). Symbionts can be further distinguished along a specialization gradient as they range from generalists that can make use of different resources to highly specialized organisms characterized by extensive trait specialization (Chomicki et al., 2020). Facultative symbionts typically do not exhibit extensive trait specialization and have a broad host range. In obligate symbionts, a more complete generalist-to-specialist specialization continuum can be observed. As specialization increases in obligate symbionts, there is often a corresponding rise in host specificity, meaning that these specialized symbionts tend to focus on a much more limited range of host species than facultative and obligate generalists.

Evolution to increasing ecological specialization and host specificity is expected to affect all aspects of symbiont biology (Bronstein, 2015; Futuyma & Moreno, 1988) and encompasses modifications in body structure and shape, the development of specialized feeding apparatus, the evolution of social behaviour, but also physiological changes (Poulin, 2011). As symbiotic animals become more specialized, their trophic and feeding ecology is predicted to undergo drastic changes (Johnson & Steiner, 2000; Nagler & Haug, 2016). Furthermore, the community of microbial symbionts in the gut or gut-associated organs of a symbiotic animal may change to better support its nutritional requirements (Kaufman et al., 2000). Indeed, bacterial symbionts are an important component of the feeding ecology of various animals, including insects (Engel & Moran, 2013; Hansen & Moran, 2014; Martens et al., 2011; Patel et al., 2014). Nutritional bacterial symbionts can be acquired vertically from parent to offspring across host generations. In insects, vertical acquisition is achieved by intracellular (transovarial) transmission but also by various extracellular transmission routes, including via specialized capsules and symbiont-supplemented jelly that covers eggs (Fukatsu & Hosokawa, 2002; Kaiwa et al., 2014; Salem et al., 2017). Stable symbioses with bacterial partners can also be mediated by the environment. For instance, environmental bacteria can postnatally colonize insect guts every generation (Kikuchi et al., 2007, 2011). In stinkbugs, this mode of environmental symbiont transmission generates complex co-infections and promiscuous bacterial symbiotic associations (Kikuchi et al., 2011). In (eu)social insects, bacterial gut symbionts can spread within a colony through social conspecific interactions (Engel & Moran, 2013; Onchuru et al., 2018). Here, coprophagy (the consumption of conspecific feces) and trophallaxis (transfer of oral fluids via mouth-to-mouth or anus-to-mouth feeding) play an important role in bacterial symbiont transmission (Koch & Schmid-Hempel, 2011; Lanan et al., 2016; Nalepa, 2015; Powell et al., 2014). In contrast, to what extent intimate heterospecific interactions between a social host and a symbiotic animal shape the microbial communities is poorly understood (but see Perry et al., 2021).

Ant nests house a surprisingly rich fauna of symbiotic arthropods known as myrmecophiles (Hölldobler & Kwapich, 2022; Parmentier, De Laender, et al., 2020). Evolution of myrmecophily independently occurred in different lineages of arthropods, and resulted in an entire continuum of trait specialization and host specificity. Trait specialization becomes most evident through the utilization of chemical and behavioural deception strategies, as well as morphological adaptations (Hölldobler & Kwapich, 2022; Komatsu et al., 2010, 2017; Parker, 2016; von Beeren et al., 2018). Higher trait specialization in myrmecophiles enables a better social integration into ant colonies, eventually reaching a point where myrmecophiles are treated as nestmates and are met with amicable behaviors, such as grooming, transport, and trophallaxis (Hölldobler & Kwapich, 2022). In contrast, generalist myrmecophiles are often poorly integrated into ant colonies, provoking aggression and avoiding direct contact with the ant host (Hölldobler & Kwapich, 2022; Parmentier et al., 2018). Nonetheless, these generalist myrmecophiles can switch more easily between various ant host species. Although a body of work has characterized the chemical, behavioural, and morphological trait modifications of specialist myrmecophiles, how increasing specialization and social integration shape the trophic and feeding ecology of myrmecophiles remains poorly understood.

The study of specialization in myrmecophiles primarily revolves around beetles (Parker, 2016; Valdivia et al., 2023; von Beeren et al., 2018), but a striking specialization continuum can also be found in silverfish, an early-branching group of wingless insects (Molero-Baltanás et al., 2017; Parmentier et al., 2022). One of the most remarkable groups is the radiated *Neoasterolepisma* genus (Zygentoma: Lepismatidae), exhibiting varying degrees of association with ants to obtain food. *Neoasterolepisma* silverfish are particularly diverse in the Iberian peninsula (Molero-Baltanás et al., 2017). Obligate generalist *Neoasterolepisma* species such as *N. curtiseta* associate with multiple ant species and tend to forage at the periphery of ant colonies, hereby avoiding direct interactions with ant workers (Molero-Baltanás et al., 2017; Parmentier et al., 2022). In contrast, another group of closely related *Neoasterolepisma* evolved specialized interactions with granivorous *Messor* ants. These myrmecophiles demonstrate the most advanced behavioural and chemical specializations within this genus and reside in the densest chambers of the nest (Parmentier et al., 2022).

Here, we studied how social integration (= specialization) modulates the feeding and trophic ecology of myrmecophiles within the *Neoasterolepisma* genus. To better understand these eco-evolutionary processes, we also studied facultative generalist and obligate generalist silverfish outside the *Neoasterolepisma* genus (of the Lepismatidae and the Nicoletiidae family, respectively). We combined stable isotope profiling, trophic tests, phylogenetic reconstruction, and microbiome characterization. We compared the isotopic niche of the generalist-specialist spectrum of ant-associated silverfish with their ant hosts using δ^15^N and δ^13^C stable isotopes. The isotopic niche, based on δ^13^C and δ^15^N values, is a proxy for trophic niche, with the δ^15^N value reflecting the trophic position and δ^13^C the basal resource in the diet of the organism (Layman et al. 2007). We performed controlled feeding experiments to further explore the diet of the myrmecophilous silverfish. Finally, we investigated the phylogenetic position of these diverse silverfish species and dissected the microbial communities of ant hosts and ant-associated silverfish.

## Materials and methods

### The silverfish study system

Here, we define “nest” as the micro-environment created by an ant colony. A nest does not only provide a home for an ant colony but also accommodates myrmecophilous silverfish. Based on lifestyle and phylogeny, we placed our sampled silverfish species in five categories; **(1) Host-specialized *Neoasterolepisma*** (Lepismatidae): these species are host-specialized in terms of high host specificity and their high level of social integration within ant colonies. Most species are associated with *Messor* and hereafter referred to as *Messor*-specialized *Neoasterolepisma* (Table 1). These species employ advanced chemical deception strategies and actively approach and interact with their host, provoke little aggression, and tolerate high densities of workers without demonstrable costs (Parmentier et al., 2022). These silverfish are typically found in large densities throughout the nest and even in the densest nest chambers (Video S1). The *Aphaenogaster*-specialized *Neoasterolepisma delator* is also integrated into the host colony, but exhibits lower densities than *Messor*-specialized species (Molero-Baltanás et al., 2017, Video S2). **(2) Obligate generalist *Neoasterolepisma*** (Lepismatidae): in this category we placed the species *N. curtiseta* that is obligately associated with ants, but is found in association with multiple unrelated ant taxa (Molero-Baltanás et al., 2017). *N. curtiseta* avoids ant interactions, typically residing in peripheral nest chambers. The presence of host workers in the same chamber leads to increased mortality when there is no refuge (Parmentier et al., 2022, Video S3). **(3) Obligate generalist nicoletiid**: this group includes members of the Nicoletiidae family *Proatelurina pseudolepisma* and *Atelura formicaria* that evolved obligate myrmecophily, and target a broad range of ants (Molero-Baltanás et al., 2017). *Atelura formicaria* is known to steal food droplets from ant workers (Janet, 1897). However, these species are less inclined to approach the host compared to host-specialized silverfish (Parmentier et al., 2022). **(4) Facultative generalist**: this category includes *Lepisma baetica* (Lepismatidae) that occurs in ant nests, but can also be found away from ants. *L. baetica* avoids interactions with ants, provokes strong aggression and, when found in ant nests, tends to occur at the periphery of the nest (Parmentier et al., 2022). **(5) Unassociated**: *Allacrotelsa kraepelini* and *Ctenolepisma ciliatum* normally do not live in ant nests, provoke a very strong aggression response, and therefore strictly avoid coming into contact with ants (Parmentier et al., 2022).

**Table 1.** Sampling efforts of our lepismatid and nicoletiid silverfish panel that covers divergent myrmecophilous lifestyles.

| Lifestyle | Subfamily | Species | Stable isotopes |  |  | Bacterial symbionts |  |  |
| --- | --- | --- | --- | --- | --- | --- | --- | --- |
|  |  |  | Specimen | Nest | Ant host | Specimen | Nest | Ant host |
| Aphaenogaster-specialized | Lepismatinae | *Neoasterolepisma delator* | 13 | 5 | 1 | 8 | 4 | 1 |
| Messor-specialized | Lepismatinae | *Neoasterolepisma balearicum* | 0 | 0 | 0 | 5 | 1 | 1 |
|  |  | *Neoasterolepisma spectabile* | 53 | 12 | 1 | 20 | 6 | 1 |
|  |  | *Neoasterolepisma foreli* | 0 | 0 | 0 | 4 | 3 | 1 |
|  |  | *Neoasterolepisma lusitanum* | 38 | 9 | 1 | 7 | 3 | 1 |
|  |  | *Neoasterolepisma balcanicum* | 0 | 0 | 0 | 4 | 1 | 1 |
|  |  | *Neoasterolepisma crassipes* | 0 | 0 | 0 | 3 | 1 | 1 |
|  |  | *Tricholepisma aureum* | 0 | 0 | 0 | 1 | 1 | 1 |
| Obligate generalist | Lepismatinae | *Neoasterolepisma curtiseta* | 19 | 7 | 2 | 14 | 4 | 4 |
|  | Atelurinae | *Proatelurina pseudolepisma* | 2 | 2 | 1 | 15 | 4 | 4 |
|  |  | *Atelura formicaria* | 0 | 0 | 0 | 13 | 5 | 5 |
| Facultative generalist | Lepismatinae | *Lepisma baetica* * | 11 | 2 | 1 | 8 | 3 | 2 |
| Unassociated | Lepismatinae | *Allacrotelsa kraepelini* * | 0 | 0 | 0 | 3 | 0 | 0 |
|  |  | *Ctenolepisma ciliatum* * | 10 | 1 | 1 | 3 | 0 | 0 |

Asterisks indicate those silverfish species for which individuals were collected that did not exhibit an association with an ant colony in the field.

### Field sampling, morphological classification, and gut dissections

Insects were sampled between September and December 2020 as well as in November 2022 across five European countries (Table S1). To avoid variation in baseline isotope values, all samples for stable isotope profiling were isolated from a single locality. The sampling site was a small (2.3 Ha) Mediterranean field in an urbanized region in Alcolea, near Córdoba (Spain), at the foot of the Sierra Morena mountain range (37.942357 N, −4.688559 E). Samples were taken within a period of a single week to minimize temporal effects on stable isotope profiling. Specimens were collected from nests using forceps and aspirators. We also collected specimens outside ant nests at this study site; i.e. the unassociated silverfish *Ctenolepisma ciliatum* and the facultative generalist *Lepisma baetica*. Specimens were kept alive in containers with moist plaster for 48 hours. This was to allow insects to empty their gut. Insects were subsequently transferred to Eppendorf tubes and killed at −21°C. During seed handling and milling, *Messor* colonies deposit husks, stalks, and seed parts in refuse dumps at the periphery of the nest (Parmentier, Gaju-Ricart, et al., 2020). This material was also collected for each sampled *Messor* nest. For feeding assays, digestive tract characterization, and amplicon sequencing, distinct localities were sampled in Belgium, Bulgaria, France, Italy, and Spain (Table S1). For amplicon sequencing, specimens were isolated using an aspirator, preserved in ∼95% ethanol, and stored at 4°C until DNA extraction. For this approach, plant material was collected from the refuge dumps of three *Messor* nests. For the dissection of the digestive tracts of silverfish, sample fixation was performed with alcoholic Bouin. Paraffin was used for inclusion with serial cuts of 4 µm.

### Stable isotope analysis

Ant, silverfish, and plant debris samples were dried at 50°C for 72 h. Ant gasters were removed to avoid contamination with the gut content. Samples were ground with a pestle and 0.2 to 1 mg for ants and silverfish, and approx. 1.5 mg for organic material of the homogenates was transferred into a tin measuring cup. We sampled silverfish individuals of each of the five lifestyle categories (Table 1). For each of the 12 *Messor* colonies (Table 2 and Table S1), we measured the isotopic signature of six individual ant workers taken along the worker size range. To measure the isotopic signature of the non-*Messor* ants, we ran a single composite sample for each colony based on a pool of five workers. We analysed the ratio of ^15^N/^14^N and ^13^C/^12^C of these samples using an element analyser isotope ratio mass spectrometer (EA-IRMS) (EA IsoLink interfaced through a ConFloIV to a delta Q; Thermo Scientific, Bremen). Measured ^15^N/^14^N and ^13^C/^12^C ratios were reported as delta (δ) values, in parts per thousand (‰). Isotopic ratios of nitrogen and carbon were normalised using USGS 89 (Porcine Collagen, for N: 6.25 ± 0.12 ‰ vs. AIR and for C: −18.13 ± 0.11 ‰ vs. VPDB). Acetanilide (for N: 19.56 ± 0.03 ‰ vs. AIR and for C: −29.50 ± 0.02 ‰ vs. VPDB), and B2155 (Caseine, for N: 5.83 ± 0.08 ‰ vs. AIR and for C: −26.98 ± 0.13 ‰ vs. VPDB) were used as quality controls.

**Table 2.** Sampling efforts of our ant panel that covers divergent ant species.

| Species | Stable isotopes |  | Bacterial symbionts |  |
| --- | --- | --- | --- | --- |
|  | Specimen | Nest | Specimen | Nest |
| *Messor barbarus* | 72 | 12 | 24 | 10 |
| *Messor ibericus (structor s.l.)* | 0 | 0 | 4 | 2 |
| *Messor* sp. | 0 | 0 | 3 | 1 |
| *Aphaenogaster senilis* | 4* | 4 | 10 | 4 |
| *Aphaenogaster gibbosa* | 1* | 1 | 0 | 0 |
| *Camponotus aethiops* | 2* | 2 | 0 | 0 |
| *Camponotus barbaricus* | 3* | 3 | 0 | 0 |
| *Camponotus cruentatus* | 0 | 0 | 2 | 1 |
| *Camponotus micans* | 1* | 1 | 0 | 0 |
| *Camponotus pilicornis* | 0 | 0 | 3 | 1 |
| *Cataglyphis velox* | 0 | 0 | 3 | 1 |
| *Iberoformica subrufa* | 0 | 0 | 3 | 1 |
| *Lasius flavus* | 0 | 0 | 2 | 1 |
| *Lasius niger* | 0 | 0 | 4 | 2 |
| *Pheidole pallidula* | 0 | 0 | 3 | 1 |
| *Tetramorium* sp. | 2* | 2 | 5 | 2 |
| *Formica fusca* | 0 | 0 | 2 | 1 |
| *Lasius emarginatus* | 0 | 0 | 2 | 1 |

\* multiple specimens pooled per sample

### Feeding assays to test trophallaxis and granivory in ant-associated silverfish

By conducting feeding assays with different food sources, we aimed to infer more insight into the feeding behaviour of the myrmecophilous silverfish. Silverfish from several nests were starved for one day before each trial (Table S1). Silverfish were individually housed in small Petri dishes (diameter 5 cm) with a moistened plaster bottom. Charcoal was added to the plaster to increase the visual contrast with the food sources. One of the following food sources was added: flour, fresh yeast, or a cut maggot of *Phaenicia sericata*. We video-recorded the behaviour of the silverfish under red light for 1 h. A food item was considered as accepted by the silverfish if we observed licking, dragging, or biting for at least 10 s. Some individuals were used in trials with different food sources, but each assay was always preceded by a starvation period of one day. Twelve feeding trials for each of the three food items (flour, yeast and a dead maggot) were conducted for the *Messor*-specialized *N. lusitanum,* and the obligate generalist nicoletiids *A. formicaria* and *P. pseudolepisma*.

Ants typically share regurgitated food droplets with nestmates through mouth-to-mouth feeding, a behavior known as trophallaxis (Meurville & LeBoeuf, 2020). Some myrmecophiles may also engage in trophallaxis and steal food droplets (Hölldobler & Kwapich, 2022). The presence of this feeding behavior in our collection of myrmecophilous silverfish was tested by offering 15 ant workers a cotton plugged vial with sugar water (20%) stained with blue colorant (E131, i.e. Patent Blue V) in a plaster filled arena (diameter 9.5 cm). After 24 h, these workers were transferred to another darkened arena (diameter 9.5 cm), and 15 starved workers of the same colony were added to promote trophallaxis. Silverfish found in the same nest were then also added and inspected after 48 h. Blue coloration of the gut was considered as an indicator that the silverfish engaged directly in trophallaxis or stole a sugar droplet during trophallaxis among workers. Blue coloration of the gut was externally visible in living specimens due to the semi-translucent cuticle. To verify whether coloration of the gut could have been missed in individuals, we dissected guts of specimen that lacked visible external coloration. These trophallaxis tests were conducted for four different species of myrmecophilous silverfish: one *Messor*-specialized silverfish, *Neoasterolepisma lusitanum* (*N =* 15 over 5 trials, host *M. barbarus*), one obligate generalist *Neoasterolepisma*, *N. curtiseta* (*N = 3,* host *Cataglyphis iberica*), and two obligate generalist nicoletiids: *Atelura formicaria* (*N* = 20 over 4 trials, host *Lasius flavus*), *Proatelurina pseudolepisma* (*N = 7*, host *L. niger*) (Table S1).

### Amplicon library preparation and sequencing

For these analyses, we also sampled silverfish species across the five lifestyle categories alongside their ant hosts (Table 1, Table 2, and Table S1). Insects were surface-sterilized through a 1 min immersion in 1% bleach, followed by two rounds of washing for 1 min in sterile water. DNA was extracted from whole specimens using the Quick-DNA Universal kit (BaseClear, the Netherlands). In parallel, DNA was extracted from three samples of plant debris that were collected from within *Messor* colonies. DNA extractions were performed in ten batches comprising a total of 187 samples, with a negative extraction control sample included in six of these batches. Sterility was ensured throughout the extraction process by working in a laminar flow hood. DNA concentrations were measured using Quant-iT™ PicoGreen™ dsDNA Assay Kit (Invitrogen) on a Synergy HTX plate reader (Biotek).

All DNA samples were used for amplicon library preparation, targeting two marker regions; a partial fragment of the insect mitochondrial cytochrome oxidase I gene (*COI*) and the V4 hypervariable region of the bacterial 16S rRNA gene. To prepare the amplicon library, a two-step PCR protocol was followed as described by (Kolasa et al., 2023). First, template-specific primers BF3-BR2 and 515F-806R, with Illumina adapter tails, amplified the *COI* and V4 16S rRNA regions, respectively (Elbrecht et al., 2019). After gel verification, SPRI bead-purified PCR products were used as templates for indexing PCR reactions. Here, Illumina adapters were completed and a unique combination of indexes was added to each library. Reaction conditions are provided in Table S2. To the first PCR reaction, we added a synthetic quantification spike-in; ∼1000 copies of linearized plasmid carrying artificial 16S rRNA amplification target Ec5502 (Tourlousse et al., 2016) to estimate the number of 16S rRNA copies per µg of DNA. Final libraries were pooled based on band brightness on agarose gels, purified with SPRI beads, and sequenced in a highly multiplexed NovaSeq 6000 SPrime 2×250 bp flow cell at the Swedish National Genomics Infrastructure facility in Stockholm.

### Analyses of amplicon sequence data

Amplicon read data of the *COI* and V4 16S rRNA marker regions were processed using a custom pipeline (https://github.com/catesval/Bioinformatic-pipelines). For each DNA sample, amplicon reads were divided based on primer sequences into two bulks that represented the two marker regions. For both bulks, quality-filtered reads were assembled into contigs using PEAR (Zhang et al., 2014). Per library, contigs were de-replicated and de-noised using VSEARCH and UNOISE2, respectively (Edgar, 2016b; Rognes et al., 2016). Chimeras were detected using USEARCH (version 11.0.667_i86linux32) (Edgar, 2010). Taxonomy of the contigs was determined using the SINTAX classifier and customized reference databases (SILVA for 16S rRNA (version SSU 138) and MIDORI (version UNIQ_GB239_CO1) for *COI*) (Edgar, 2016a). Contigs were clustered at the 97% identity level via the UPARSE-OTU algorithm of USEARCH (Edgar, 2010). The V4 16S rRNA zOTU data was de-contaminated using six DNA extraction blanks and three PCR blanks, based on the relative zOTU abundances across our experimental and control samples (https://github.com/catesval/Bioinformatic-pipelines). Specifically, a V4 zOTU was not considered a contaminant when the maximum relative abundance in at least one experimental sample was tenfold higher than its maximum relative abundance in any blank sample. We further designated V4 zOTUs as symbiotic when the maximum relative abundance in at least one experimental sample was higher than 0.001. Using the copy number / read number ratio of the Ec5502 spike-in, an estimate of symbiont density was obtained for each sample by calculating the number of 16S rRNA copies per ng DNA. To control for the dense populations of intracellular *Anaplasmataceae* symbionts (including *Wolbachia*), corresponding reads were first subtracted. *COI* barcoding was unsuccessful for three silverfish samples and for the three plant debris samples. For all other samples, the most abundant eukaryotic *COI* sequence variant was identified and used for further analyses.

### Statistical analyses

All analyses were carried out using R (version 4.2.2) (R Core Team, 2021). Raw data are publicly accessible at XXX and NCBI. We plotted the isotopic position of silverfish relative to their ant host workers in C-N biplots for each silverfish species separately. We grouped *Messor-*specialized *N. spectabile* and *N. lusitanum* as *Messor*-specialized *Neoasterolepisma* to increase the statistical power of our analyses. Given that both species shared nearly identical niches when co-habiting, and can only be distinguished by subtle morphological traits, we think this is an acceptable approach. We calculated a distance matrix with pairwise Euclidean distance for the *Neoasterolepisma* lifestyles, i.e. *Messor*-specialized *Neoasterolepisma*, the *Aphaenogaster-*specialized *N. delator*, and the obligate generalist *N. curtiseta*, between the samples in the C-N biplot. Using a PERMANOVA test (adonis function vegan, 999 permutations), we aimed to explain the variation in this distance matrix by the predictors taxon identity (ant or silverfish), ant nest, and their interaction. To compare the isotopic niches of the *Messor-*specialized *Neoasterolepisma* and their ant hosts, we drew nest-specific ellipses encompassing approximately 95% of the data (function plotSiberObject in SIBER). We also tested whether δ^15^N was significantly different between *Messor*-specialized *Neoasterolepisma* and their host, using a linear model with taxon (ant or silverfish) as predictor and a random slope allowing an interaction between nest origin (12 levels) and taxon (lmer). The analysis was repeated with δ^13^C as the dependent variable.

To understand the determinants of the composition and density of the bacterial community of myrmecophilous silverfish, we first tested whether these features were different among three key myrmecophilous lifestyles within the *Neoasterolepisma* genus, i.e. *Aphaenogaster*- and *Messor*-specialized and obligate generalist species. Here, we also did not account for the species identity of different *Messor*-specialized *Neoasterolepisma* species. Data was curated by filtering zOTUs using a mean read count threshold of 20 and a minimum variance threshold of 10 % (based on the standard deviation). The microbial community composition was compared based on the relative abundance of the nine most abundant bacterial orders in our silverfish panel (hereby excluding the ‘unassigned’ and ‘Other’ categories). When different individuals of the same silverfish lifestyle were sampled in a nest, we took their average microbial composition to avoid pseudo-replication (nest-specific averages). Associations between the silverfish microbial composition (Bray Curtis dissimilarity matrix) and the predictor silverfish lifestyle (three levels) were examined using a PERMANOVA test (adonis function vegan, 999 permutations). P-values of the Post hoc comparisons between two levels of silverfish lifestyles were adjusted by Benjamini Hochberg (BH) correction. Next, we tested the effect of silverfish lifestyle (three levels) on the relative abundance of Lactobacillales, the most abundant bacterial order in *Neoasterolepisma*, using non-parametric Kruskal-Wallis tests coupled with Post hoc Dunn tests with BH correction. In addition, we compared the total microbiome density across the three myrmecophilous lifestyles in *Neoasterolepisma.* Density values were log transformed and modelled against the fixed factor “lifestyle” and the random factor “nest” using a general linear mixed model. Pairwise post hoc comparisons were controlled by BH-correction.

After investigating the silverfish microbiome on the order level, we focused on the genotypic (or zOTU) diversity within the *Weissella* genus, a prominent component of the *Leuconostocaceae* family (Lactobacillales) in our system. Specifically, we tested whether the strain composition of *Weissella* bacteria of *Messor-*specialized *Neoasterolepisma* silverfish and *Messor* ants was determined by nest origin or by taxon (ant or silverfish). The *Weissella* strain profile was based on the presence/absence of the *Weissella* zOTUs that were retained after data curation. We performed a PERMANOVA analysis (adonis function vegan, 999 permutations) on the Jaccard distance matrix of the *Weissella* strain profiles. This analysis assesses the impact of the predictors taxon (ant or silverfish), nest origin, and their interaction by partitioning the explained variation among these predictors.

### Phylogenetic reconstruction

For all phylogenetic analyses, the consensus *COI* nucleotide sequences were aligned by the E-INS-i strategy of MAFFT, leaving gappy regions (Katoh & Toh, 2010). Models were selected based on the Bayesian Information Criterion using ModelFinder (Kalyaanamoorthy et al., 2017). Maximum-likelihood tree reconstructions and ultrafast bootstrapping (5,000 replicates) were performed with IQ-TREE (version 2.1.2) (Hoang et al., 2018; Minh et al., 2020) (random seed number was set at 54321). UFBoot trees were optimized by nearest neighbour interchange (argument ‘-bnni’).

## Results

We sampled a total of 58 ant colonies for workers and ant-associated silverfish and sampled four ant-free localities for unassociated silverfish (Table 1, Table 2, and Table S1). In addition, we also sampled plant debris material produced by *Messor* workers during seed handling and milling. To study the stable isotope profiles of our insect panel, we analyzed 243 samples. In total, bacterial 16S rRNA and mitochondrial *COI* amplicon data were generated and analyzed for 178 individual insects and three plant debris samples. Based on the curated *COI* amplicon data and conservative morphological classification, our insect panel consisted of 18 ant species and 14 silverfish species (Table 1 and Table 2).

### Host specialized silverfish display a corresponding isotopic niche with their ant host

Based on the stable isotope analyses, the ant-silverfish food web showed high trophic differentiation in our sample collection (Figure 1). The δ^13^C of our study species ranged from - 28.2 to −23.5 ‰ (a range of δ^13^C values reflects a variability of basal food sources) and δ^15^N from 3.2 to 7.8 ‰ (higher δ^15^N values reflects higher trophic positions) (Figure 1). The averaged isotopic signatures of the *Messor*-specialized silverfish *N. lusitanum* and *N. spectabile* closely grouped together with those of their *Messor* host (Figure 1). Three silverfish species (*N. curtiseta*, *P. pseudolepisma*, and *N. delator*) occupied the highest trophic positions on average. However, only the obligate generalist nicoletiid *P. pseudolepisma* and the obligate generalist *Neoasterolepisma* (*N. curtiseta*) displayed an average enrichment in ^15^N relative to their ant host. The *Aphaenogaster-*specialized *N. delator* also grouped closely with its host in the δ^13^C-δ^15^N biplot (Figure 1). *Lepisma baetica* (unassociated and associated with *Tetramorium*) and *C. ciliatum* were characterized by average δ^13^C and δ^15^N values in our insect panel. The different ant hosts could be clearly differentiated in the isotopic biplot. *Camponotus* occupied on average the lowest trophic position, while *Messor* and *Tetramorium* had on average an intermediate trophic position. *Camponotus* ants were further characterized by the highest levels of δ^13^C.

**Figure 1.**
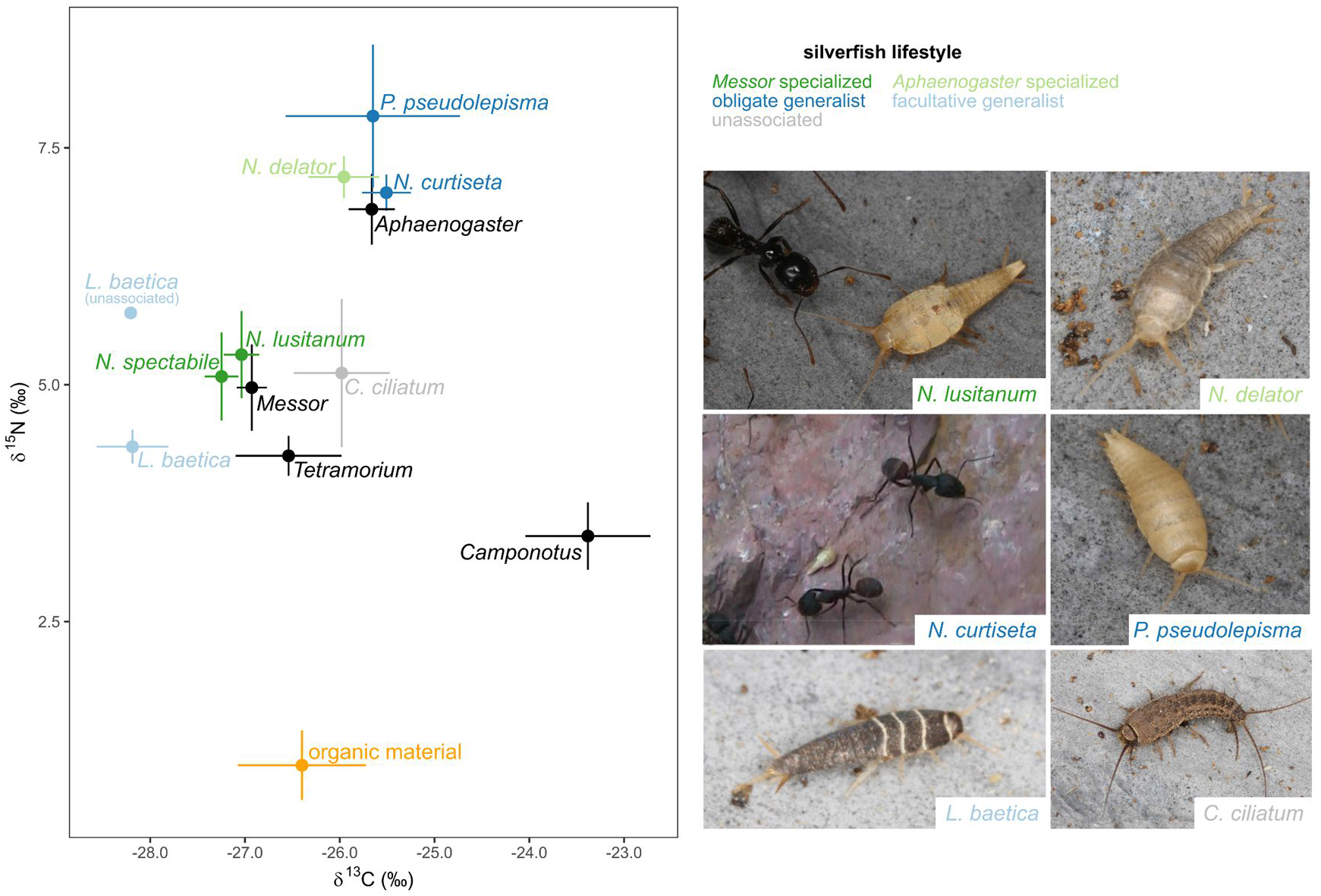
The average isotopic niche of different silverfish and ant species varies greatly within a single sampling locality. When multiple specimens of the same species were sampled within a nest, we calculated the average specific to that particular nest. Consequently, each data point represents the mean value of these nest-specific averages for the species (e.g., *N. spectabile* based on 12 nest averages of 53 individuals in total, see Table 1). Error bars correspond to standard errors (SE). Colours represent different lifestyles and sample types; dark green: *Messor*-specialized *Neoasterolepisma* (Video S1), light green: *Aphaenogaster*-specialized *Neoasterolepisma* (Video S2), dark blue: obligate generalist *Neoasterolepisma* (*N. curtiseta,* Video S3) and nicoletiid (*P. pseudolepisma*), light blue: facultative generalist *Lepisma baetica*, light grey: unassociated, black: ant hosts, and orange: organic material of *Messor* nests. The mean values of ant-associated and unassociated *Lepisma baetica* individuals are given separately. Silverfish belonging to the different lifestyle categories are illustrated in the right panel. Caption color corresponds to the lifestyle group.

*Messor* workers occupied an intermediate trophic position averaged across the 12 colonies (Figure 1). However, when focusing on individual nests, a strong intercolonial variation of isotopic niches in the C-N biplot was observed, especially for δ^15^N-values (Figure 2a, b). Certain *Messor* colonies exhibited exceedingly high δ^15^N values, a finding that appears at odds with its granivorous lifestyle. The difference in δ^15^N values between the colony with the highest (δ^15^N = 8.5 ± 0.2) and the lowest trophic position (δ^15^N = 2.9 ± 0.2) was 5.6 (Figure 2a,b). For both *Messor-*specialized *Neoasterolepisma* species (*N*. *lusitanum* and *N*. *spectabile*), there was a strong match in the isotopic niche with the co-habiting *Messor* workers (Figure 2a,b, Figure S1). No differences in either δ^15^N (lmer, ꭓ² = 2.1, df = 1, *P* = 0.14) or δ^13^C (ꭓ² = 2.8, df = 1, *P* = 0.10) were found between *Messor-*specialized silverfish and their *Messor* host. The isotopic niches of *Messor*-specialized silverfish showed medium to strong overlap in 10 out of 12 nests to those of the *Messor* hosts (average overlap of 42.5%, range: 28.1 - 89.0%, Figure S1). Note the overlap between the two species of *Messor-specialized Neoasterolepisma* (Figure S1). The isotopic distance (Euclidean distance in the CN-biplot) between these *Messor*-specialized silverfish and their host workers (living in the same nest environment) was much shorter compared to the distance to ant workers from other *Messor* nests (ratio: 2.7 ± 0.2 SE). This clustering at the nest level of *Messor* ants and *Messor*-specialized silverfish was also supported by a PERMANOVA approach. The R^2^-values in this analysis give the proportion of explained variance by each of the predictor variables. Most variation in the isotopic niche of these silverfish and their *Messor* hosts was explained by their shared nest environment (PERMANOVA, R² = 0.84, *P =* 0.001), rather than by their specific taxon identity (PERMANOVA, R² = 0.02, *P =* 0.001) or the interaction between taxon and nest (PERMANOVA, R² = 0.04, *P =* 0.001) (Figure 2a, b, Figure S1). In parallel, the isotopic niche of the *Aphaenogaster-*specialized silverfish *N. delator* and their *Aphaenogaster* host was also mainly driven by the nest environment (PERMANOVA, *N. delator-Aphaenogaster*: R² = 0.57, *P =* 0.08; distance to *Aphaenogaster* workers found in the same nest was 3.6 ± 1.8 SE times smaller than to *Aphaenogaster* workers from other nests in the isotope biplot) (Figure 2c). Taxon identity (R² = 0.03, *P =* 0.52) and the interaction with nest (R² = 0.08, *P =* 0.78) explained a small fraction of the observed variation in isotopic niche of the *Aphaenogaster-*specialized *N. delator* and *Aphaenogaster* ants. The obligate generalist *Neoasterolepisma* (*N. curtiseta*), the obligate generalist nicoletiid *P. pseudolepisma* and the facultative generalist *L. baetica* had a larger isotopic distance to their hosts than the *Messor-* and *Aphaenogaster*-specialized *Neoasterolepisma* species (Figure 2d-f). The distance between the obligate generalist *N. curtiseta* and co-habiting workers and non-co-habiting *Camponotus* workers was similar (ratio 1.0 ± 0.1 SE). Isotopic niche variation in the δ^13^C-δ^15^N biplot of *N. curtiseta* and its *Camponotus* hosts was mainly determined by taxon identity (R² = 0.62, *P =* 0.001) and to a lesser degree to nest environment (R² = 0.23, *P =* 0.004) or the interaction of nest with taxon (R² = 0.06, *P =* 0.17) (Figure 2e).

**Figure 2.**
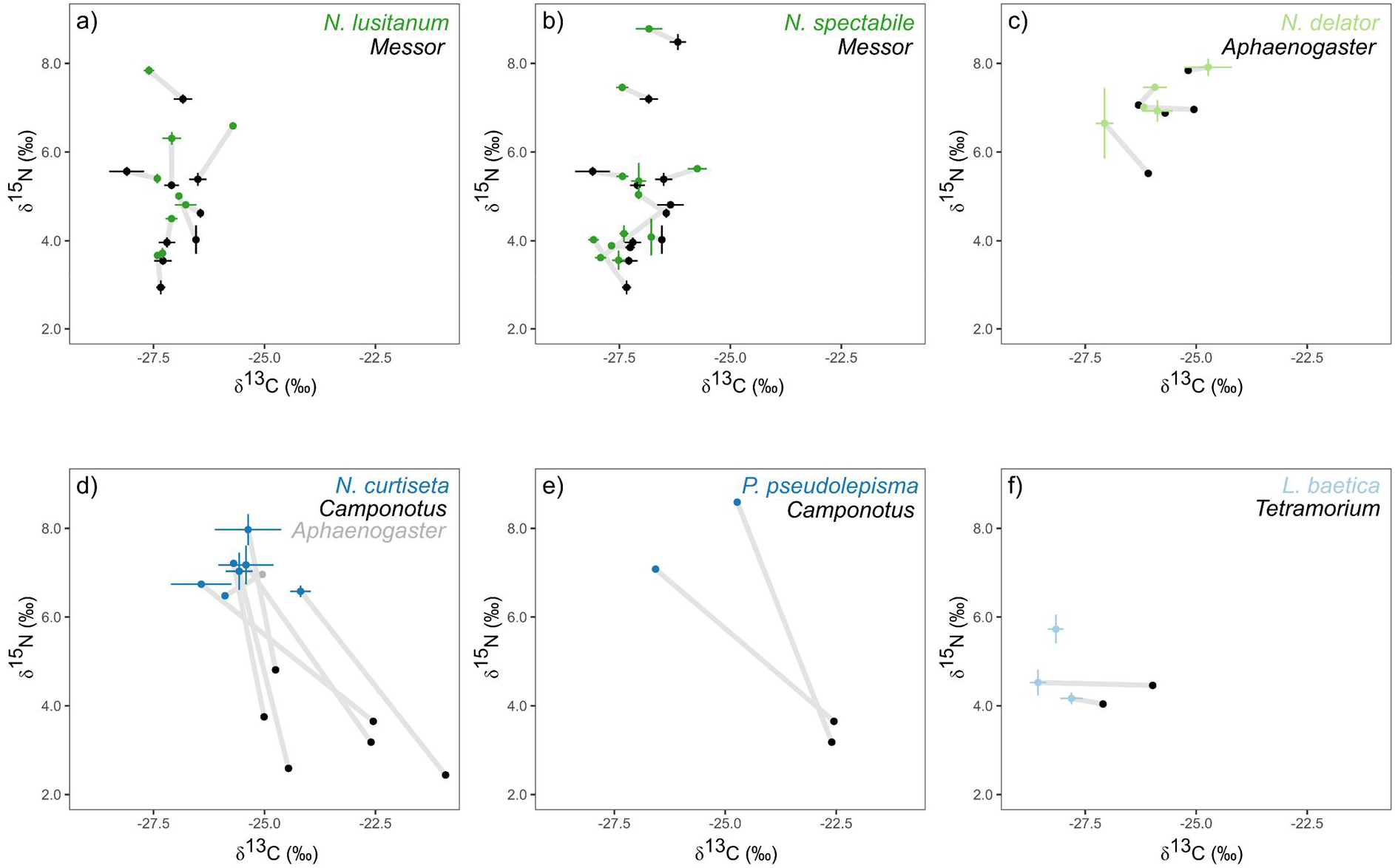
The isotopic niche converges between host-specialized silverfish and ant hosts inhabiting the same nest. Different panels give the variation of δ^13^C and δ^15^N of silverfish species and their ant hosts across different nests. Each point represents the average ant worker (black or gray) or silverfish value (coloured) per nest: a) *Messor* and *Messor-* specialized *N. lusitanum*, b) *Messor* and *Messor-*specialized *N. spectabile*, c) *Aphaenogaster* and *Aphaenogaster-*specialized *N. delator*, d) obligate generalist nicoletiid *P. pseudolepisma* and *Camponotus* ants, e) obligate generalist *N. curtiseta* and *Camponotus* and *Aphaenogaster* ants, f) facultative generalist *L. baetica* and *Tetramorium* host. Error bars correspond to standard errors (SE). A gray band connects ant workers and silverfish from the same nest.

### Feeding behaviour and physiology of ant-associated silverfish

Flour, yeast, and cut maggots were consistently neglected in the feeding assays with the starved obligate generalist nicoletiids *P. pseudolepisma* and *A. formicaria*. The *Messor-* specialized *N. lusitanum* also ignored yeast and dead maggots, but was observed feeding on flour in three of the twelve conducted trials (Video S4). The gut of six out of 20 *A. formicaria* individuals and three out of 7 *P. pseudolepisma* individuals was clearly coloured, indicating the acquisition of food droplets from their ant host through mouth-to-mouth feeding. This confirmed previous observations of this behavior in the generalist nicoletiid *A. formicaria* (Janet, 1897) (Figure S2). The mouth parts of some *P. pseudolepisma* individuals were also stained, further supportive of heterospecific trophallaxis (Figure S2). When dissecting the gut of one specimen of *P. pseudolepisma* without visible external body colouration, a light blue staining of the gut was still observed. In contrast, no evidence of trophallaxis was detected in the obligate generalist *N. curtiseta* and *Messor*-specialized *N. lusitanum*.

To further study the feeding modes of our panel of silverfish, we dissected the gut of *N. curtiseta*, *N. spectabile*, and *P. pseudolepisma.* Consistent with previous observations of Lepismatidae (Barnhart, 1961; Pothula et al., 2019), *N. curtiseta* and *N. spectabile* displayed a differentiated proventriculus that carries sclerotized teeth-like structures (Figure S3). These findings show that the evolution of myrmecophily within *Neoasterolepisma* silverfish is not associated with a secondary loss of this feeding structure, indicating that ant-associated *Neoasterolepisma* likely still rely on solid food for nutrition. In contrast, dissections of *P. pseudolepisma* specimens did not reveal a muscular proventriculus. Moreover, the digestive tract of *P. pseudolepisma* lacked sclerotized teeth-like structures (Figure S4). These findings are in line with previous studies comparing the digestive system of silverfish families (Wygodzinsky, 1961) and align with the hypothesis that trophallaxis could be an important mode of feeding in *P. pseudolepisma* and *A. formicaria* (Janet, 1897).

### Myrmecophilous lifestyle determines the composition and density of the silverfish microbiome

Morphological classification of the insect specimens was collaborated by the mitochondrial *COI* amplicon data. For the myrmecophiles, however, we did observe identical *COI* sequences for *Tricholepisma aureum* and *Neoasterolepisma balearicum*, raising the possibility of potential synonymous taxonomic descriptions (Figure 3, Figure S6, and Data S1). Hereafter, we included *T. aureum* within the *Neoasterolepisma-*specific analyses and considered its lifestyle as *Messor-*specialized *Neoasterolepisma*. In addition, based on the *COI* data, the silverfish species *P. pseudolepisma* and *A. formicaria* appeared to consist of cryptic genetic lineages (Figure 3 and Data S1). The mitochondrial *COI* phylogeny placed our *Messor* collection into four species; *Messor barbarus*, *Messor ibericus (structor s.l*.), *Messor structor s.l*. (tentatively morphologically classified as *Messor mcarthuri* by Lech Borowiec) (Steiner et al., 2018), and one unidentified *Messor* species, hereafter referred to as *Messor* sp. (Figure S7).

**Figure 3.**
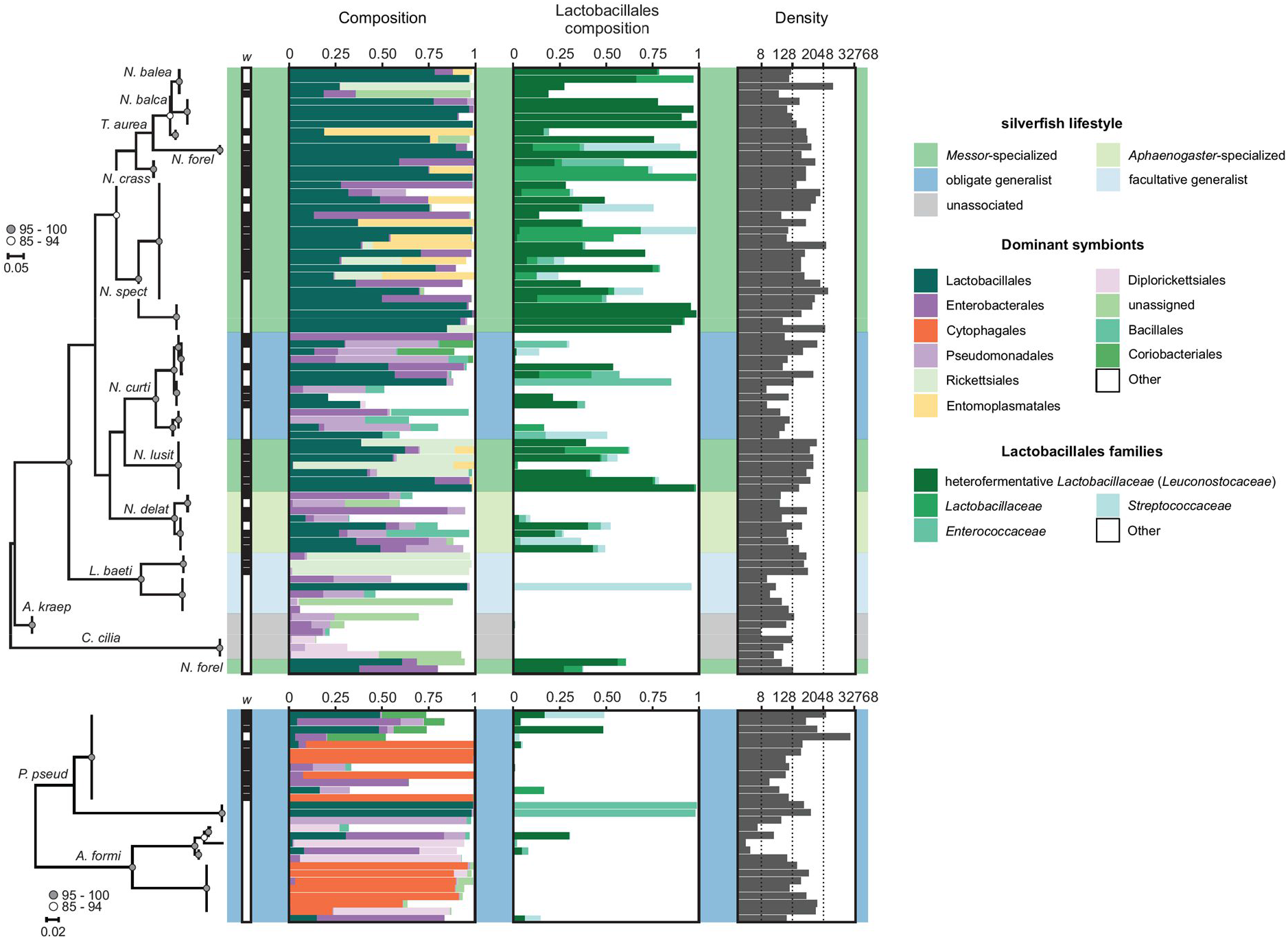
Myrmecophilous lifestyle shapes the silverfish microbiome. The Lepismatinae and Atelurinae phylogenies are depicted and are based on a 418 bp fragment of the mitochondrial *COI* gene. *COI* amplification was not successful for three silverfish samples and their symbiotic communities are depicted below their respective phylogenetic trees. Silverfish lifestyle is colour-coded (see right). *Wolbachia* infection is indicated by a black background within the row labelled with ‘*w*’. The unit for the microbiome density is the number of symbiotic rRNA copies per ng DNA. *Leuconostocaceae* is currently considered as a later synonym of *Lactobacillaceae*.

Insects are often colonized by dense populations of maternally transmitted bacterial endosymbionts that manipulate host reproduction (Perlmutter & Bordenstein, 2020). Amplicon sequence data revealed that of these reproductive endosymbionts, *Wolbachia* (of the *Anaplasmataceae* family) were the most common in our insect panel, infecting ∼36 % of the individuals (Figure 3, Figure S5, Figure S6, and Figure S7). *Wolbachia* variants were clustered into four OTUs and 18 zOTUs, representing supergroups A, B, and F (Figure S5). *Wolbachia* typically reach high densities within host reproductive tissues and are maternally transmitted across host generations (Kaur et al., 2021; Perlmutter & Bordenstein, 2020). In our dataset, we generally observed different 16S rRNA genotypes in different species (Figure S5), further arguing against recent horizontal transmission of *Wolbachia* (Figure 3, Figure S5, Figure S6, and Figure S7). To focus on bacterial (gut) symbionts that may horizontally transmit, we removed amplicon reads belonging to the intracellular *Anaplasmataceae* family.

Excluding the ‘unassigned’ and ‘Other’ categories, the nine most common bacterial orders comprised on average 81 % of the silverfish microbiome (Figure 3 and Data S1). Lactobacillales was the most dominant order and was represented by four families; heterofermentative *Lactobacillaceae* (*Leuconostocaceae*), *Streptococcaceae*, *Lactobacillaceae*, and *Enterococcaceae* (Figure 3). The taxonomic designation *Leuconostocaceae* is now considered to be synonymous with *Lactobacillaceae* and, for sake of clarity, we will refer to this clade as heterofermentative *Lactobacillaceae* hereafter (Zheng et al., 2020). Heterofermentative *Lactobacillaceae* symbionts commonly infected *Neoasterolepisma* silverfish and reached high relative abundance levels (Figure 3). In contrast, the three Lepismatinae silverfish species outside the *Neoasterolepisma* genus (*L. baetica*, *C. ciliatum*, and *A. kraepelini*) did not carry heterofermentative *Lactobacillaceae* (Figure 3). Obligate generalist nicoletiids *P. pseudolepisma* and *A. formicaria* did carry these lactic acid bacterial symbionts, albeit at a lower prevalence (Figure 3). For Enterobacterales, *Enterobacter* bacteria were the most common. Within *Neoasterolepisma* silverfish, infections of several vertically transmitted symbionts were inferred, including *Spiroplasma* (Entomoplasmatales) and *Rickettsia* (Rickettsiales) (Figure 3). All three specimens of unassociated *C. ciliatum* carried a *Rickettsiella* infection (Diplorickettsiales), as did some *A. formicaria* insects. Within our silverfish panel, intracellular *Cardinium* symbionts (Cytophagales) were restricted to 12 specimens of the two obligate generalist nicoletiids (*P. pseudolepisma* and *A. formicaria*) where high relative abundance levels were observed, representing maternally acquired infections (Figure 3 and Data S1) (Duron et al., 2008).

Within the *Neoasterolepisma* silverfish genus, the bacterial composition (based on the nine most abundant bacterial orders) was significantly determined by the myrmecophilous lifestyle (PERMANOVA, R² = 0.39, F = 5.81, *P* = 0.001). Post-hoc comparisons revealed that the bacterial composition was significantly different in *Messor*-specialized *Neoasterolepisma* compared to obligate generalist *Neoasterolepisma* (Post hoc PERMANOVA, R² = 0.24, F = 4.8, BH corrected *P =* 0.018), and *Aphaenogaster-*specialized *Neoasterolepisma* (Post hoc PERMANOVA, R² = 0.38, F = 9.23, BH corrected *P =* 0.003). The relative difference in microbial composition in *Messor-*specialized *Neoasterolepisma* compared to the two other lifestyles was mainly driven by a higher proportion of Lactobacillales (Kruskal-Wallis test, *P =* 0.005, mean of the nest-averaged proportions ± SE: *Messor*-specialized = 0.62 ± 0.06, *Aphaenogaster*-specialized = 0.14 ± 0.10, and obligate generalist = 0.32 ± 0.09). Post hoc Dunn tests revealed that the microbiome of *Messor-*specialized *Neoasterolepisma* had a higher proportion of Lactobacillales compared to *Aphaenogaster*-specialized *Neoasterolepisma* (BH corrected *P =* 0.008), and a near-significant higher proportion than obligate generalist *Neoasterolepisma* (BH corrected *P =* 0.083).

In addition, the myrmecophilous lifestyle also significantly determined the gut bacterial density within *Neoasterolepisma* silverfish (lmer, ꭓ² = 11.1, df = 2, *P* = 0.004) (Figure 3). Comparisons uncovered that gut bacterial densities were significantly higher for *Messor-* specialized *Neoasterolepisma* compared to the obligate generalist *N. curtiseta* (*P =* 0.04) and to *Aphaenogaster*-specialized *N. delator* (*P* = 0.04) (Figure 3).

To gain a first insight into how myrmecophily modifies the microbiome of *Neoasterolepisma* silverfish, we also investigated the symbiotic community of the ant hosts (Figure S7 and Data S1). In ants, Lactobacillales was also among the most dominant orders and was represented by three families; heterofermentative *Lactobacillaceae* (*Leuconostocaceae*), *Streptococcaceae*, and *Lactobacillaceae* (Figure S7). Of our *Messor* collection, ∼71 % of the workers exhibited an infection of heterofermentative *Lactobacillaceae* symbionts (Figure S7). For *Aphaenogaster*, three workers (30 %) carried an apparent infection (Figure S7). Of the ant hosts of the obligate generalist *N. curtiseta*, only a single specimen of *Pheidole pallidula* appeared infected with heterofermentative *Lactobacillaceae*. *Camponotus* ants were infected with beneficial *Blochmannia* endosymbionts (Enterobacterales), in line with a rich body of literature (de Souza et al., 2007; Feldhaar et al., 2007). In contrast to our silverfish panel, all other sampled ant species exhibited a relatively low infection prevalence of vertically transmitted symbionts.

### Nest explains the *Weissella* infection profile of *Messor*-specialized *Neoasterolepisma* silverfish

To further dissect the origins and modes of transmission of the heterofermentative *Lactobacillaceae* symbionts, we investigated the repertoire of genetic lineages within this bacterial family. Here, we also included samples of organic seed remnants of three *Messor* nests. A total of four OTUs and 22 zOTUs remained after data curation (Figure 4, Figure S8, and Figure S9). OTU4 was assigned with 97 % certainty to the heterofermentative *Lactobacillaceae* clade but could not be confidently placed on the genus level, and is hereafter referred to as Lacto4. Lacto4 was observed in ∼77 % of all *Messor-*specialized *Neoasterolepisma* silverfish. In contrast, the *Aphaenogaster-*specialized and obligate generalist *Neoasterolepisma* silverfish (*N. delator* and *N. curtiseta*, respectively) were not infected with Lacto4, nor were any other silverfish outside the *Neoasterolepisma* genus (Figure 4, Figure S8, and Figure S9). Of the ant collection, only two *Messor* workers (∼7 %) appeared positive for Lacto4 (Figure 4, Figure S8, and Figure S9). None of the seed samples exhibited the presence of Lacto4 bacteria. Together, these findings strongly suggest that Lacto4 infection is associated with the myrmecophilous lifestyle of *Messor-*specialized *Neoasterolepisma*.

**Figure 4.**
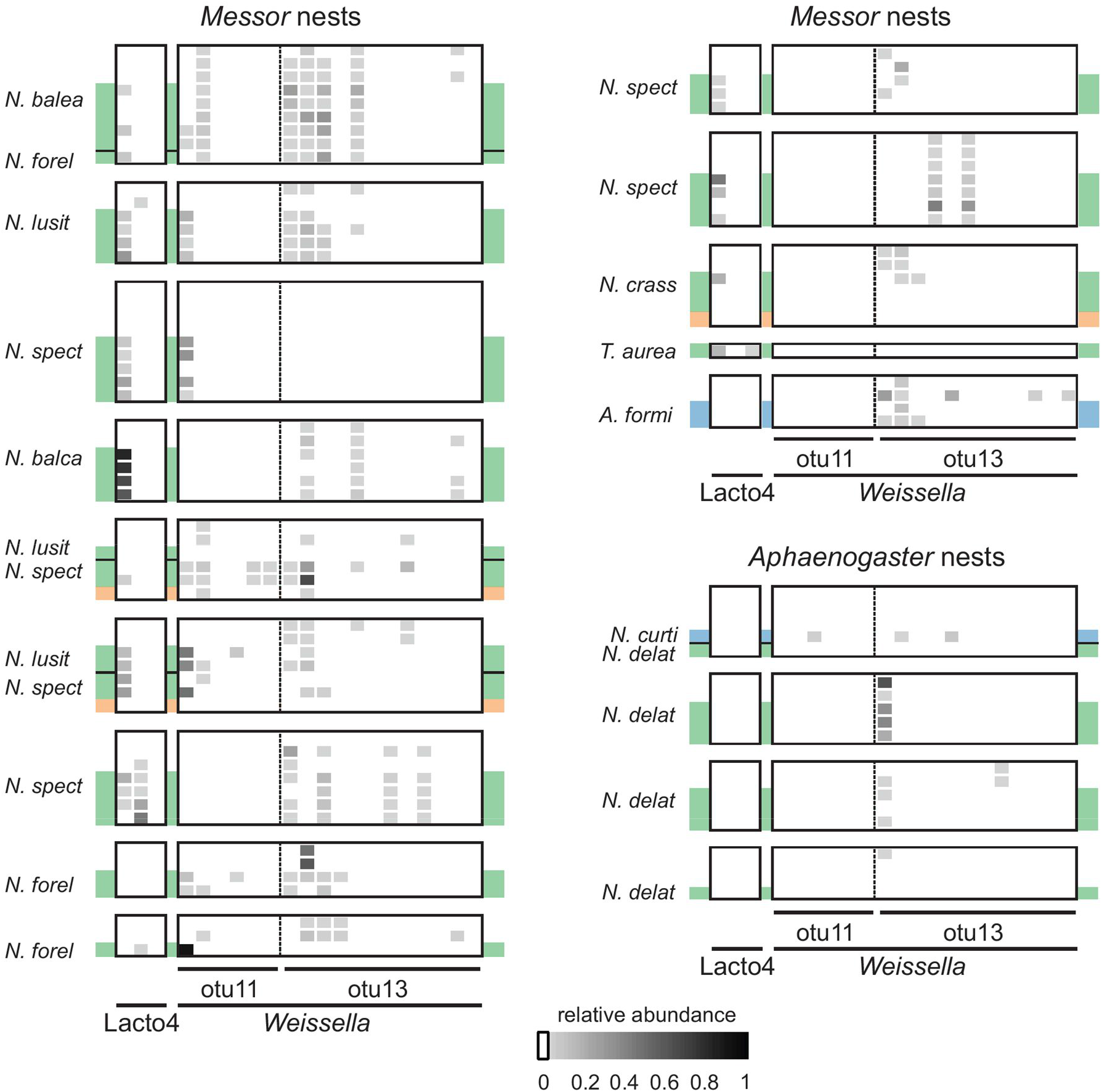
Ant nest determines the infection profile of *Weissella* bacteria in *Messor*-specialized silverfish. Ant and silverfish individuals are grouped according to nest origin. The relative abundances of heterofermentative *Lactobacillaceae* symbionts are visualized (see bottom). Lacto4 and *Weissella* bacteria depict apparently distinct transmission modes within this system. Silverfish lifestyle is colour-coded; dark green: *Messor-*specialized *Neoasterolepisma*, light green: *Aphaenogaster-*specialized, and blue: obligate generalist (*N. curtiseta* and *A. formicaria*). Plant seed material is indicated with an orange background. Silverfish species are identified by their abbreviated species name.

Within the heterofermentative *Lactobacillaceae* clade, we also observed two distinct lineages (i.e. OTUs) of *Weissella* that infected our silverfish and ant specimens (Figure 4 and Data S1). Here, *Weissella* exhibit a strong association with *Messor-*specialized silverfish and *Messor* ants (Figure 4, Figure S8, and Figure S9). We identified 6 and 12 strains (i.e. zOTUs) for OTU11 and OTU13, respectively. In most nests, we observed a strong overlap in the composition of these 18 *Weissella* strains between silverfish and ants (Figure 4). This heterospecific sharing of *Weissella* strains in *Messor* nests was statistically confirmed by variation partitioning using a PERMANOVA approach. Variation in the composition of these 18 strains was mostly explained by the nest origin of the insect (PERMANOVA, R² = 0.63, F = 11.89, *P* = 0.001). Taxon (ant or silverfish) and the interaction between taxon and nest also significantly impacted the *Weissella* infection profile (PERMANOVA, taxon, R² = 0.04, F = 7.33, *P* = 0.001, taxon:nest, R² = 0.15, F = 2.82, *P* = 0.001), but predicted a lower proportion of the variation in *Weissella* composition than nest origin. One OTU of the heterofermentative *Lactobacillaceae* clade, OTU26 (associated with a single zOTU, zOTU38), was only found in a single *C. pilicornis* colony where *P. pseudolepisma* and *N. curtiseta* individuals tested positive (Figure S9).

## Discussion

Specialization shapes the ecology and evolution of symbiotic animals. In symbiotic arthropods living with eusocial insects, chemical and behavioral specialization enable them to access the innermost chambers of the host nest without triggering aggression. This social integration into the colony enables symbiotic arthropods to optimally benefit from the favorable microclimate and resources inside a well-defended nest fortress (Hölldobler & Kwapich, 2022). Compared to specialization strategies as body morphology, behavior and chemical deception, relatively little is known about the relationship between social integration (specialization) and feeding ecology. In this study, we demonstrate that the convergence of feeding ecology of socially integrated silverfish and their ant hosts may be an important aspect of their ecology and evolution.

The isotopic niches of *Messor*- and *Aphaenogaster*-specialized *Neoasterolepisma* exhibited a strong similarity to those of their host colonies in the isotope biplot. This tight relationship implies that these silverfish not only feed on similar food sources but also flexibly adapt their diet in accordance with the foraging decisions of the host colony. Previously, coupling of the isotopic niche among co-habiting animals have been shown in host-parasite systems (Gómez-Díaz & González-Solís, 2010), co-habiting ant species (Sprenger et al., 2021), co-habiting termite species (Florencio et al., 2013) and myrmecophiles and their ant host (Parmentier et al., 2023). In these studies, although the δ^13^C-δ^15^N (or only δ^15^N) profiles of the co-habiting species covary, there was a clear difference in the isotopic composition of the co-habiting species. Here, we show that the isotopic niches of co-habiting animals do not only covary but can also converge to the same position in C-N isotopic space. *Messor* ants are generally considered as granivorous and are consequently expected to have a relatively low trophic position, just above the seed material that they consume (Platner et al., 2012). Nonetheless, some *Messor* colonies of our study displayed significantly enriched δ^15^N values, exceeding those of the organic seed remnants in their waste dumps by more than one trophic level (when considering a standard enrichment of 3‰-4‰ per trophic position, Post 2002). Relatively enriched δ^15^N values in *Messor* colonies were previously found by Fiedler *et al*. (2007), leading the authors to suggest that the reliance on resources other than plant seed may be greater in *Messor* than commonly believed (Fiedler et al., 2007). Fiedler *et al*. (2007) further hypothesized that the enriched δ^15^N values observed in *Messor* could be attributed to a regular consumption of arthropod remains, animal excrement, or fungi growing on organic nest material. Enriched δ^15^N values may also reflect small-scale spatial variability in the baseline, but we did not find any spatial autocorrelation in δ^15^N values of the organic plant material in the waste dumps or of the ant species. Lastly, seeds of different plants may strongly vary in δ^15^N (Wissinger et al., 2014) and the variation in trophic position across colonies might reflect the preferential consumption of seeds with distinct δ^15^N values. But this explanation seems implausible as we did not detect a correlation with the δ^15^N of the organic seed remains in the waste deposits. In strong support of our study, *Messor*-specialized *Neoasterolepisma* attained similarly enriched levels of δ^15^N in those *Messor* colonies with unusually high δ^15^N. Feeding trials under controlled laboratory conditions showed that *Messor*-specialized *Neoasterolepisma* feed on plant material just like their granivorous *Messor* hosts. Based on our results, it is unlikely that dead ants or ant brood form a significant part of the diet of *Messor*-specialized *Neoasterolepisma*, otherwise one would expect a notable enrichment in ^15^N compared to their *Messor* host (Parmentier et al., 2016). *Messor*-specialized *Neoasterolepisma* also do not engage in trophallaxis (unlike obligate generalist nicoletiid species). A recent study underlined the importance of the diet quality of a termite host for the fitness of an associated beetle (Eloi et al., 2023). *Messor*-specialized silverfish are also subject to the quality of the food resources consumed by their ant hosts, but it is unknown whether these silverfish select nests based on the nest feeding ecology. *Aphaenogaster* ants are considered as omnivorous (Seifert, 2007) and had on average a higher trophic position than *Messor* ants. Like the *Messor*-specialized species, we found a strong convergence in the isotopic niche between *Aphaenogaster*-specialized silverfish and *Aphaenogaster* ants.

In sharp contrast, congeneric *N. curtiseta* silverfish did not show a resemblance of their isotopic niche compared to their ant hosts but showed a stronger enrichment in δ^15^N. This obligate generalist has a broad host range, but was mainly found with *Camponotus* ants across the study sites. The low position of our *Camponotus* samples is consistent with other studies (Blüthgen et al., 2003; Fiedler et al., 2007). The obligate generalist *N. curtiseta* is less integrated into their host colonies than *Messor*- and *Aphaenogaster*-specialized *Neoasterolepisma* (Parmentier et al., 2022). While *Messor* and *Aphaenogaster* specialists are expected to feed on food resources within the nest, *N. curtiseta* may be less restricted to the nest environment and scavenge for food sources outside the nest. Note that the trophic position of *N. curtiseta* was higher than the unassociated silverfish *Ctenolepisma*, which is thought to have a standard diet for silverfish.

The myrmecophilous lifestyle of *Neoasterolepisma* was also associated with dramatic changes in the bacterial microbiome. Heterofermentative lactic acid bacteria were significantly associated with *Messor*-specialized *Neoasterolepisma,* and the infection patterns suggest two distinct modes of transmission for Lacto4 and *Weissella* symbionts within this silverfish system. As Lacto4 symbionts only displayed a high relative abundance in *Messor-*specialized *Neoasterolepisma*, we hypothesize that Lacto4 symbionts might (mainly) spread via vertical transmission in these silverfish. Currently, the most parsimonious scenario is that an ancestral *Messor*-specialized *Neoasterolepisma* species acquired Lacto4 before derived lineages formed the contemporary set of *Messor*-specialized species. Unfortunately, the current study lacks the phylogenetic depth for both *Neoasterolepisma* and Lacto4 to formally test whether Lacto4 indeed vertically transmits within this system, and when the acquisition of Lacto4 occurred. Radiations of insect species with new feeding ecologies have previously been coupled to the acquisition of vertically transmitted bacterial symbionts. Herbivory is typically considered as a key innovation and many insect taxa engage in symbiosis with bacteria to survive on a herbivorous diet. In tortoise leaf beetles, bacterial *Stammera* symbionts encode pectinases that allow its beetle host to digest pectin-rich foliage (Salem *et al*. 2017, 2020).

Sap-feeding insects such as spittlebugs and cicadas depend on a diverse array of nutritional symbionts for the provision of essential amino acids and vitamins (Moran *et al*. 2008). Here, we are drawn to the hypothesis that the acquisition of Lacto4 symbionts facilitated the shift towards a (semi-)granivorous feeding ecology of *Messor-*specialized *Neoasterolepisma.* Bacteria of this clade are reported to contribute to the degradation and modification of recalcitrant plant material, including complex polysaccharides (Siezen et al., 2006). The investigation of whether Lacto4 may contribute to the assimilation of plant nutrients in *Messor-* specialized *Neoasterolepisma* is an interesting avenue of future research for this system.

We also observed *Weissella* symbionts of the heterofermentative lactic acid bacteria clade in our insect collection. *Weissella* have been isolated from various plants and animals, including insects (Berasategui et al., 2016; Heo et al., 2019; Oh et al., 2013; Praet et al., 2015). Here, two distinct lineages of *Weissella* co-infect silverfish and ants. Moreover, the *Weissella* strain profile of *Messor*-specialized silverfish and *Messor* ants converges and is strongly determined by the nest, suggestive of two non-mutually exclusive transmission scenarios. First*, Weissella* bacteria may initially infect plant seeds and subsequently colonize silverfish and ant guts upon feeding. Studies have shown that the microbiotic communities of insect herbivores can be contingent on the microbiome of the host plant (Chung et al., 2017; Mogouong et al., 2021; Pirttilä et al., 2023; Strano et al., 2018). For our focal system, this scenario implies that *Weissella* acquisition is likely associated with the granivorous feeding behaviour of *Messor*-specialized *Neoasterolepisma.* The observation that a seed sample from a *Messor* colony tested positive for both *Weissella* lineages supports this transmission scenario. Second, *Weissella* infection could be further maintained by horizontal transmission between silverfish and ants within nests. The behaviour of *Messor*-specialized *Neoasterolepisma* to engage in close physical contact with *Messor* workers likely creates multiple opportunities for heterospecific transmission. Here, our laboratory assays show that trophallaxis or grooming might not be essential behavioural traits to allow for heterospecific transmission of bacteria within ant nests. Unfortunately, the current data does not allow to reliably identify the original source of *Weissella,* nor the direction of potential horizontal transmission routes. *Weissella* have also been detected in other myrmecophile-ant systems. Based on amplicon and Sanger read data, *Weissella* infection was apparently shared across myrmecophilous beetles and ants within colonies of the army ant *Eciton burchellii* (Valdivia *et al*., 2023). However, no effect of ant colony on the *Weissella* infection profile was readily apparent. Instead, infection was associated with the development of *E. burchellii* with larvae carrying a relatively high abundance of *Weissella.* Further study is required to fully understand *Weissella* transmission and potential functional roles in insects.

Using the silverfish model system, we uncovered that social hosts strongly determine various features of the feeding ecology of symbiotic animals. We show that the isotopic niches of co-habiting animals can converge to the same position in C-N isotopic space. We further uncovered that the social environment can explain the changes in the density and composition of the microbiome of symbiotic animals.

## Electronic supplementary material

Table S1. Sampling details of the insect panel.

Table S2. PCR reaction conditions.

Figure S1. Nest-specific isotopic niche overlap between *Messor* host and *Messor-*specialized *Neoasterolepisma*.

Figure S2. The generalist nicoletiids *P. pseudolepisma* and *A. formicaria* steal food droplets from ant hosts.

Figure S3. *Neoasterolepisma* silverfish display a differentiated proventriculus with sclerotized teeth-like structures.

Figure S4. The nicoletiid *P. pseudolepisma* lacks sclerotized teeth-like structures in its digestive tract.

Figure S5. Maternally transmitted *Wolbachia* bacteria are pervasive in our insect panel.

Figure S6. Myrmecophilous lifestyle shapes the silverfish microbiome (with insect specimen ID).

Figure S7. Heterofermentative *Lactobacillaceae* symbionts infect *Messor* ants.

Figure S8. Ant nest determines the infection profile of *Weissella* bacteria in *Messor*-specialized silverfish.

Figure S9. Heterofermentative *Lactobacillaceae* symbionts are generally restricted to host*-* specialized *Neoasterolepisma* within our silverfish panel.

Video S1. *Messor*-specialized *Neoasterolepisma* in an opened *Messor* nest.

Video S2. *Aphaenogaster*-specialized *Neoasterolepisma* (*N. delator*) running across a stone that covered their *Aphaenogaster* host nest.

Video S3. Generalist *Neoasterolepisma* (*N. curtiseta*) running across a stone that covered their *Camponotus* host nest.

Video S4. *Messor*-specialized *Neoasterolepisma* feeding on flour.

Data S1. Microbiome OTU and zOTU data.

## Acknowledgements

We thank Alberto Tinaut for his help in the classification of some of our ant specimens and Albena Lapeva-Gjonova for kindly providing insect samples. We are indebted to Katja Van Nieuland and Elise Verstraete for their assistance with the EA-IRMS experiments, and to Mateusz Buczek for assistance with amplicon sequencing library preparation. NW was supported by a Special Research Fund (BOF) postdoctoral fellowship (Ghent University, 01P03420), TP by a Research Foundation – Flanders (FWO, 1203020N) and a Fund for Scientific Research postdoctoral fellowship (FNRS, 30257865), PŁ by the Polish National Agency for Academic Exchange (NAWA) grant PPN/PPO/2018/1/00015, and CV by the Polish National Science Centre (NCN) grant 2018/31/B/NZ8/01158.

## Author Contributions

TP and NW conceived and designed the experiments. TP, RMB, MGR, and NW performed the experiments. PB and PŁ contributed essential data, infrastructure, and reagents. TP, RMB, CV, MGR, and NW analyzed the data. TP and NW wrote the manuscript with input from all authors.

## Statement on inclusion

Our study brings together authors from a number of different countries, including scientists based in the countries where the study was carried out. All authors were engaged early on with the research and study design to ensure that the diverse sets of perspectives they represent were considered from the onset. Whenever relevant, literature published by scientists from the region was cited.

