## Supporting info for "Co-habiting ants and silverfish display a converging feeding ecology"

### Table S1. Sampling details of the insect panel

| Nest/Location | Analysis | Coordinates/City | Country | Ant species | Silverfish species |
|---|---|---|---|---|---|
| N1 | M | 36°56’07’’ N, 3°23’21’’ W | Spain | *Cataglypis velox* | *Neoasterolepisma curtiseta* |
| N2 | M | 36°56’07’’ N, 3°23’21’’ W | Spain | *Messor barbarus* | *Neoasterolepisma spectabile* |
| N3 | M | 36°56’07’’ N, 3°23’21’’ W | Spain | *Messor barbarus* | *Neoasterolepisma spectabile* |
| N4 | M | 37°09’20’’ N, 4°09’21’’ W | Spain | *Iberoformica subrufa* | *Neoasterolepisma curtiseta* |
| N5 | M | 37°09’20’’ N, 4°09’21’’ W | Spain | *Messor barbarus* | *Neoasterolepisma foreli* |
| N6 | M | 37°09’20’’ N, 4°09’21’’ W | Spain | *Messor barbarus* | *Neoasterolepisma foreli* |
| N7 | M | 37°09’20’’ N, 4°09’21’’ W | Spain | *Messor barbarus* | *Neoasterolepisma spectabile* |
| N8 | M | 37°09’20’’ N, 4°09’21’’ W | Spain | *Messor* sp. | *Neoasterolepisma spectabile* |
| N9 | M | Córdoba | Spain | *Camponotus pilicornis* | *Neoasterolepisma curtiseta*<br>*Lepisma baetica*<br>*Proatelurina pseudolepisma* |
| N10 | M | Córdoba | Spain | *Aphaenogaster senilis* | *Neoasterolepisma curtiseta*<br>*Neoasterolepisma delator* |
| N11 | M | Córdoba | Spain | *Pheidole pallidula* | *Proatelurina pseudolepisma* |
| N12 | M | Córdoba | Spain | *Lasius niger* | *Proatelurina pseudolepisma* |
| N13 | M | Córdoba | Spain | *Aphaenogaster senilis* | *Neoasterolepisma delator* |
| N14 | M | Córdoba | Spain | *Aphaenogaster senilis* | *Neoasterolepisma delator* |
| N15 | M | Córdoba | Spain | *Tetramorium* sp. | *Lepisma baetica* |
| N16 | M | Córdoba | Spain | *Tetramorium* sp. | *Lepisma baetica* |
| N17 | M | 41°29’34’’ N, 2°00’49’’ E | Spain | *Messor barbarus* | *Neoasterolepisma balearicum*<br>*Tricholepisma aureum*<br>*Neoasterolepisma foreli* |
| N18 | M | Córdoba | Spain | *Aphaenogaster senilis* | *Neoasterolepisma delator* |
| N19 | M | Córdoba | Spain | *Messor barbarus* | *Neoasterolepisma lusitanum* |
| N20 | M | Golyam Dervent | Bulgaria | *Messor ibericus* (*structor s.l.*) | *Neoasterolepisma balcanicum* |
| N21* | M | Córdoba | Spain | *Messor barbarus* | *Neoasterolepisma spectabile* |
| N22* | M | Córdoba | Spain | *Messor barbarus* | *Neoasterolepisma spectabile*<br>*Neoasterolepisma lusitanum* |
| N23 | M | 45°34’40.4” N, 10°46’37.0” E | Italy | *Messor ibericus* (*structor s.l.*) | *Atelura formicaria* |
| N24 | M | 45°34’41.0” N, 10°46’45.5” E | Italy | *Lasius emarginatus* | *Atelura formicaria* |
| N25 | M | 45°34’36.2” N, 10°46’51.8” E | Italy | *Formica fusca* | *Atelura formicaria* |
| N26* | M | 43°50'4.0"N, 3°44'29.6"E | France | *Messor barbarus* | *Neoasterolepisma crassipes* |
| N27 | M | 43°50'2.2"N, 3°44'30.2"E | France | *Camponotus cruentatus* | *Proatelurina pseudolepisma* |
| N28 | M | 43°50'12.3"N, 3°44'28.6"E | France | *Messor* sp. | *Tricholepisma aureum* |
| N29 | M/F | 51°10'33.3" N, 2°47'41.4" E | Belgium | *Lasius flavus* | *Atelura formicaria* |
| N30 | M | 51°10'32.7" N, 2°47'42.2" E | Belgium | *Lasius niger* | *Atelura formicaria* |
| M1 | SI | 37°56’32.2” N, 4°41’16.0” W | Spain | *Messor barbarus* | *Neoasterolepisma spectabile*<br>*Neoasterolepisma lusitanum* |
| M2 | SI | 37°56’33.5” N, 4°41’16.1” W | Spain | *Messor barbarus* | *Neoasterolepisma spectabile*<br>*Neoasterolepisma lusitanum* |
| M3 | SI | 37°56’33.3” N, 4°41’16.4” W | Spain | *Messor barbarus* | *Neoasterolepisma spectabile* |
| M4 | SI | 37°56’32.5” N, 4°41’15.8” W | Spain | *Messor barbarus* | *Neoasterolepisma spectabile*<br>*Neoasterolepisma lusitanum* |
| M5 | SI | 37°56’32.8” N, 4°41’16.3” W | Spain | *Messor barbarus* | *Neoasterolepisma spectabile*<br>*Neoasterolepisma lusitanum* |
| M6 | SI | 37°56’31.9” N, 4°41’19.2” W | Spain | *Messor barbarus* | *Neoasterolepisma spectabile* |
| M7 | SI | 37°56’31.3” N, 4°41’21.9” W | Spain | *Messor barbarus* | *Neoasterolepisma spectabile*<br>*Neoasterolepisma lusitanum* |
| M8 | SI | 37°56’31.6” N, 4°41’23” W | Spain | *Messor barbarus* | *Neoasterolepisma spectabile*<br>*Neoasterolepisma lusitanum* |
| M9 | SI | 37°56'31.4" N, 4°41'21.9" W | Spain | *Messor barbarus* | *Neoasterolepisma spectabile*<br>*Neoasterolepisma lusitanum* |
| M10 | SI | 37°56'31.7" N, 4°41'22.6" W | Spain | *Messor barbarus* | *Neoasterolepisma spectabile*<br>*Neoasterolepisma lusitanum* |
| M11 | SI | 37°56’31.9” N, 4°41’25.2” W | Spain | *Messor barbarus* | *Neoasterolepisma spectabile*<br>*Neoasterolepisma lusitanum* |
| M12 | SI | 37°56’31.2” N, 4°41’26.3” W | Spain | *Messor barbarus* | *Neoasterolepisma spectabile* |
| A1 | SI | 37°56’33.3” N, 4°41’16.6” W | Spain | *Aphaenogaster senilis* | *Neoasterolepisma delator* |
| A2 | SI | 37°56’43.6” N, 4°40’40.1” W | Spain | *Aphaenogaster senilis* | *Neoasterolepisma delator* |
| A3 | SI | 37°56’33.3” N, 4°41’15.9” W | Spain | *Aphaenogaster gibbosa* | *Neoasterolepisma delator* |
| A4 | SI | 37°56’33.2” N, 4°41’16.0” W | Spain | *Aphaenogaster senilis* | *Neoasterolepisma delator* |
| A7 | SI | 37°56’31.7” N, 4°41’28.1” W | Spain | *Aphaenogaster senilis* | *Neoasterolepisma delator*<br>*Neoasterolepisma curtiseta* |
| C1 | SI | 37°56’33.3” N, 4°41’16.7” W | Spain | *Camponotus aethiops* | *Neoasterolepisma curtiseta* |
| C2 | SI | 37°56’32.2” N, 4°41’20.4” W | Spain | *Camponotus aethiops* | *Neoasterolepisma curtiseta* |
| C3 | SI | 37°56’32.1” N, 4°41’20.8” W | Spain | *Camponotus micans* | *Neoasterolepisma curtiseta*<br>*Ctenolepisma ciliatum* |
| C4 | SI | 37°56’33.6” N, 4°41’16.2” W | Spain | *Camponotus barbaricus* | *Neoasterolepisma curtiseta*<br>*Proatelurina pseudolepisma* |
| C6 | SI | 37°56’30.3” N, 4°41’26.2” W | Spain | *Camponotus barbaricus* | *Neoasterolepisma curtiseta* |
| C7 | SI | 37°56’31.7” N, 4°41’25.4” W | Spain | *Camponotus barbaricus* | *Neoasterolepisma curtiseta*<br>*Proatelurina pseudolepisma* |
| T1 | SI | 37°56’31.8” N, 4°41’22.7” W | Spain | *Tetramorium* group *semilaeve* | *Lepisma baetica* |
| T2 | SI | 37°56’31.8” N, 4°41’22.7” W | Spain | *Tetramorium* group *forte* | *Lepisma baetica* |
| X1 | M | | Spain | unasssociated | *Allacrotelsa kraepelini* |
| X2 | M | | Spain | unasssociated | *Ctenolepisma ciliate* |
| X3 | SI | 37°56'31.7"N 4°41'20.1"W | Spain | unasssociated | *Lepisma baetica* |
| X4 | SI | 37°56'32.1"N 4°41'21.0"W | Spain | unasssociated | *Ctenolepisma ciliate* |
| F1 | F | 38°54'25.6"N 4°47'39.3"W | Spain | *Messor barbarus* | *Neoasterolepisma lusitanum* |
| F2 | F | 38º24'11" N, 5º25'44" W | Spain | *Cataglyphis iberica* | *Neoasterolepisma curtiseta* |
| F3 | F | 37º55'23" N, 4º48'09" W | Spain | *Lasius niger* | *Proatelurina pseudolepisma* |

An asterisk indicates those *Messor* nests where organic plant material was sampled for microbiome analysis. For stable isotope profiling, organic plant material of each *Messor* nest was collected. The column ‘Analysis’ indicates whether individuals were collected for microbiome characterization (M), stable isotope profiling (SI), or feeding trials (F).

**Table S2. PCR reaction conditions**

| Step | Temp [ºC] | | Time [s] | | Cycles | |
|---|---|---|---|---|---|---|
| | PCR1 | PCR2 | PCR1 | PCR2 | PCR1 | PCR2 |
| Initial denaturation | 95 | 95 | 900 | 900 | 1 | 1 |
| Denaturation | 94 | 94 | 30 | 30 | 25 | 7 |
| Annealing | 50 | 50 | 90 | 90 | 25 | 7 |
| Extension | 72 | 72 | 90 | 90 | 25 | 7 |
| Final extension | 72 | 72 | 600 | 600 | 1 | 1 |
| Store | 8 | 8 | ∞ | ∞ | | |

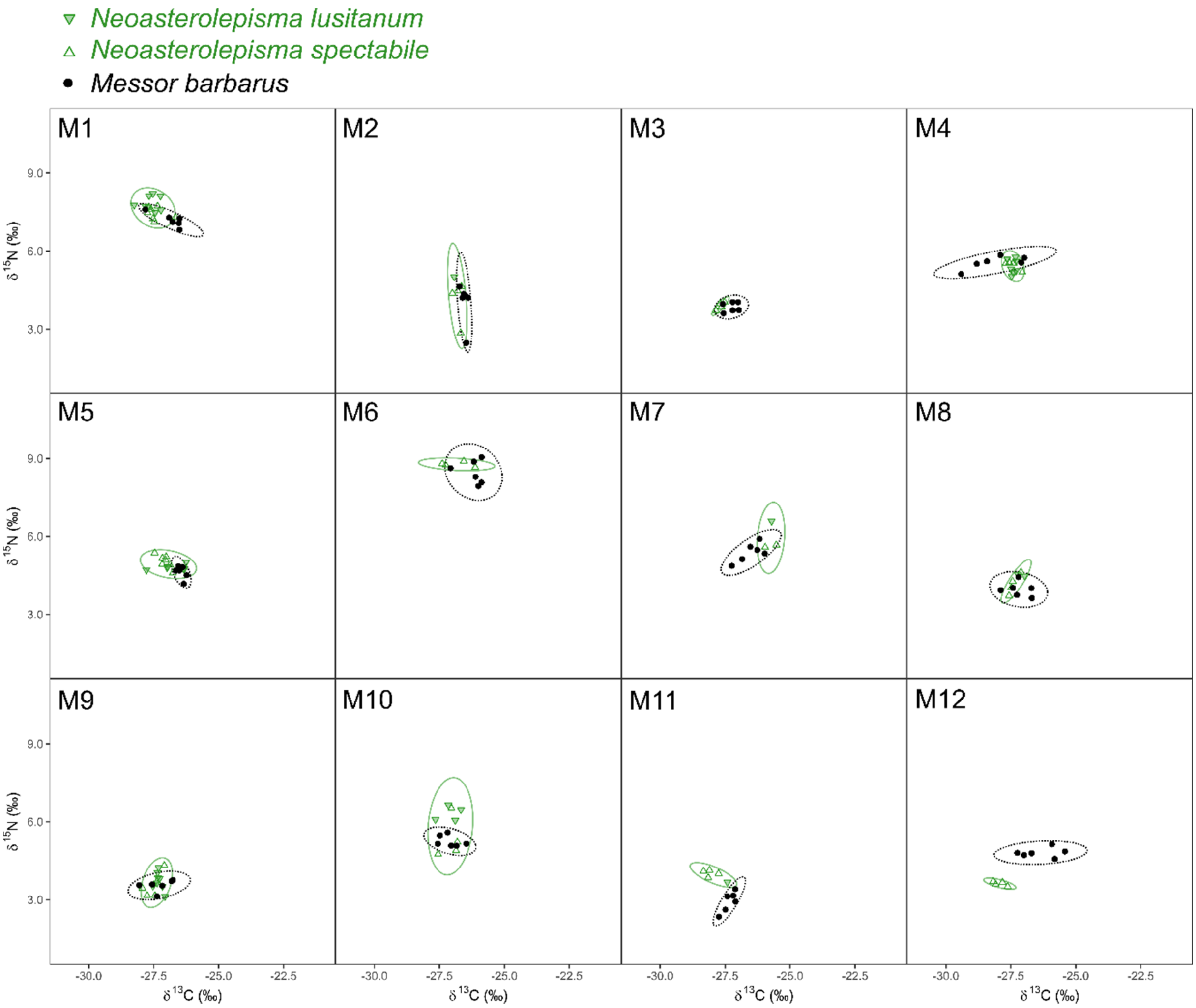


**Figure S1. Nest-specific isotopic niche overlap between *Messor* host and *Messor*-specialized *Neoasterolepisma*.** Each point represents the $\delta^{13}C$-$\delta^{15}N$ signature of an individual *M. barbarus* worker (black dot) or *Messor*-specialized *Neoasterolepisma* (down-pointing green triangle *N. lusitanum*, up-pointing green triangle *N. spectabile*). The *M. barbarus* workers and silverfish are grouped per nest, with each of the twelve panels corresponding to one of the twelve sampled *M. barbarus* nests.: M1, M2, …, M12. Note that both *Messor*-specialized species co-occurred in multiple nests. Ellipses comprise approx. 95% of the *Messor* worker samples (black ellipse) or *Messor*-specialized *Neoasterolepisma* (green ellipse).

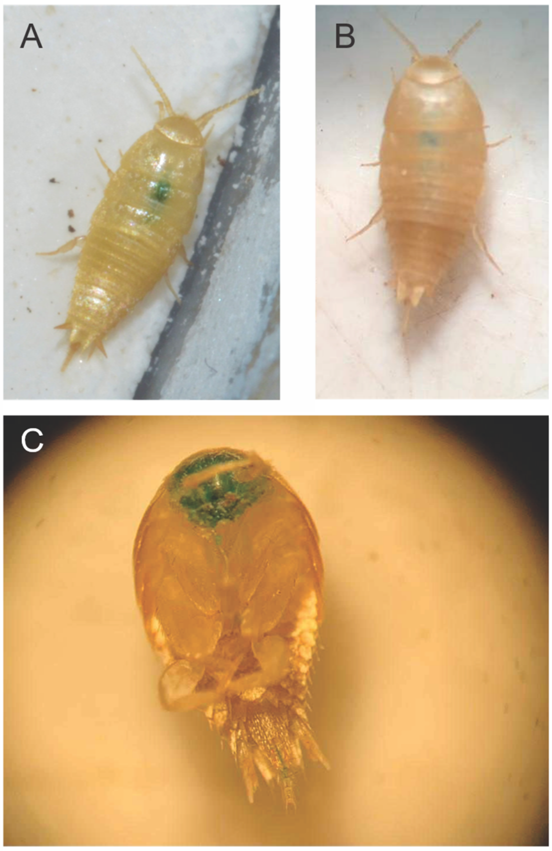


**Figure S2. The obligate generalist nicoletiids *P. pseudolepisma* and *A. formicaria* steal food droplets from ant hosts. A:** Our trophallaxis assays confirm previous observations of this behaviour in *A. formicaria* (Janet, 1897). **B:** Stealing of food droplets was also observed for the obligate generalist nicoletiid *P. pseudolepisma.* **C:** The mouth parts of a number of *P. pseudolepisma* individuals were stained.

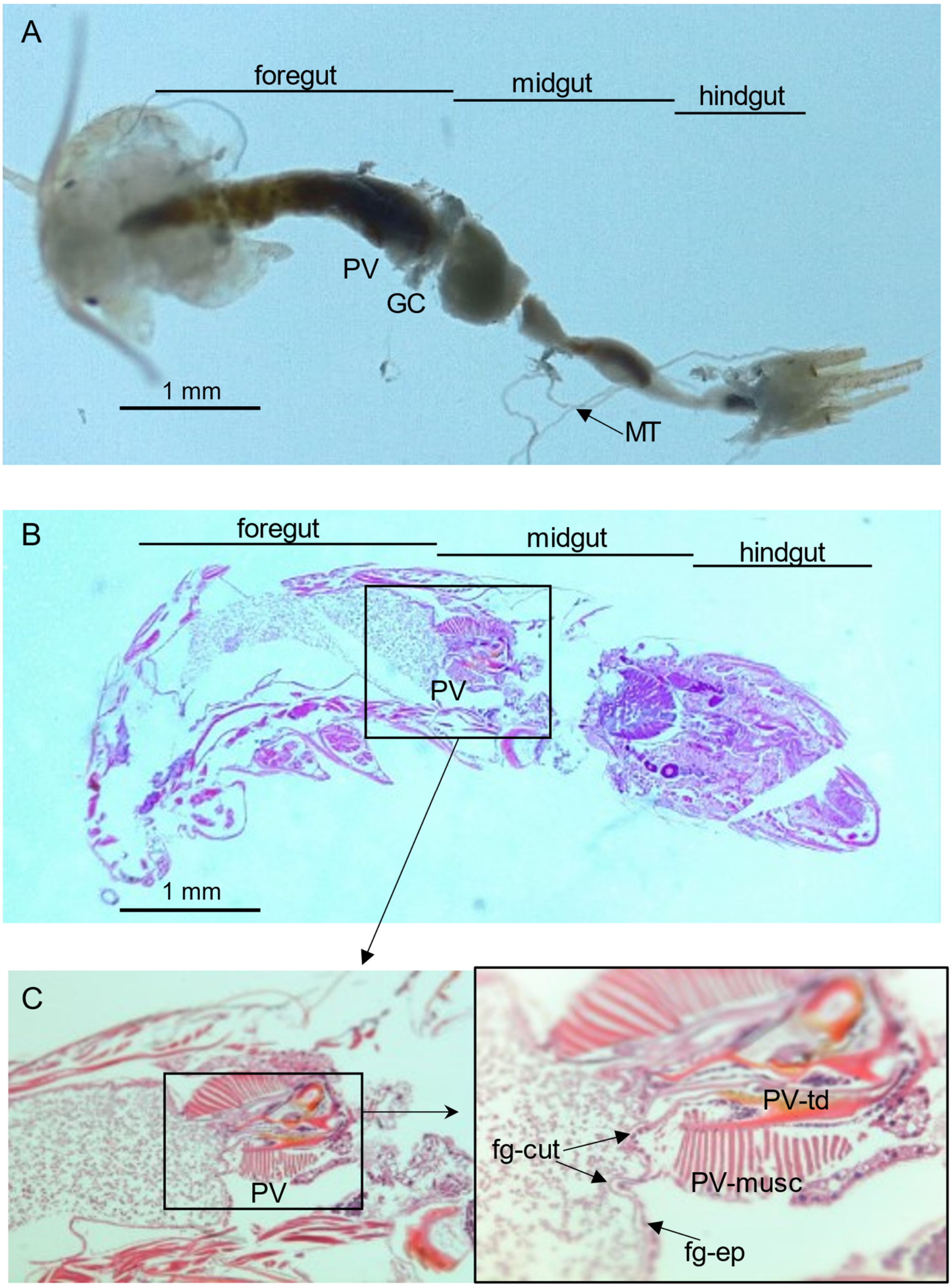


**Figure S3. *Neoasterolepisma* silverfish display a differentiated proventriculus with sclerotized teeth-like structures.** Throughout panels A to C, abbreviations are; fg-cut: foregut cuticle, fg-ep: foregut epithelium, GC: gastric caeca, MT: Malpighian tubules, PV-musc: proventriculus musculature, PV-td: proventriculus tooth-like denticles, SV: stomodeal valve. A Masson trichromic stain was used.

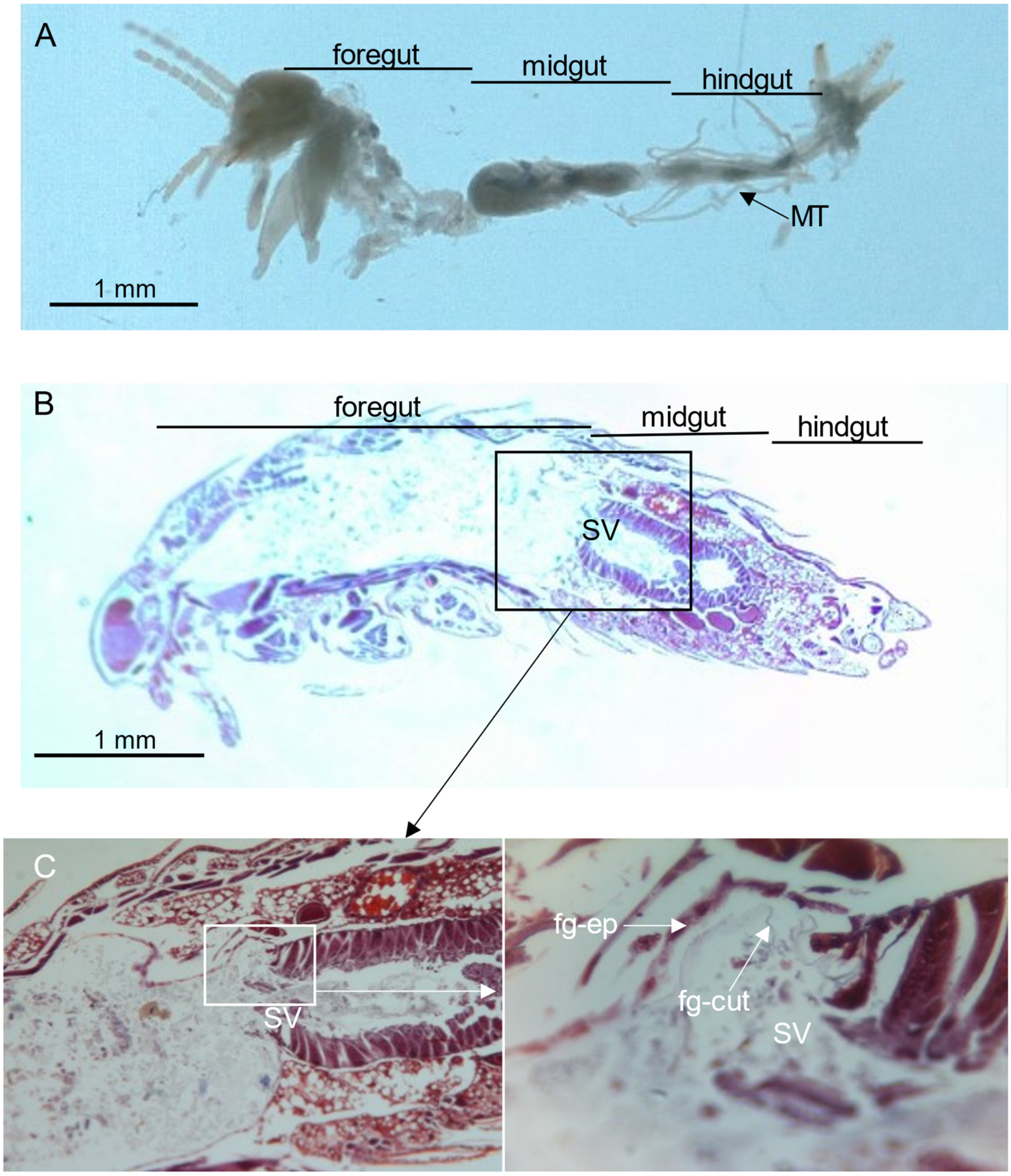


**Figure S4. The nicoletiid *P. pseudolepisma* lacks sclerotized teeth-like structures in its digestive tract.** Throughout panels A to C, abbreviations are; fg-cut: foregut cuticle, fg-ep: foregut epithelium, MT: Malpighian tubules, and SV: stomodeal valve. Hematoxylin eosin stain was used.

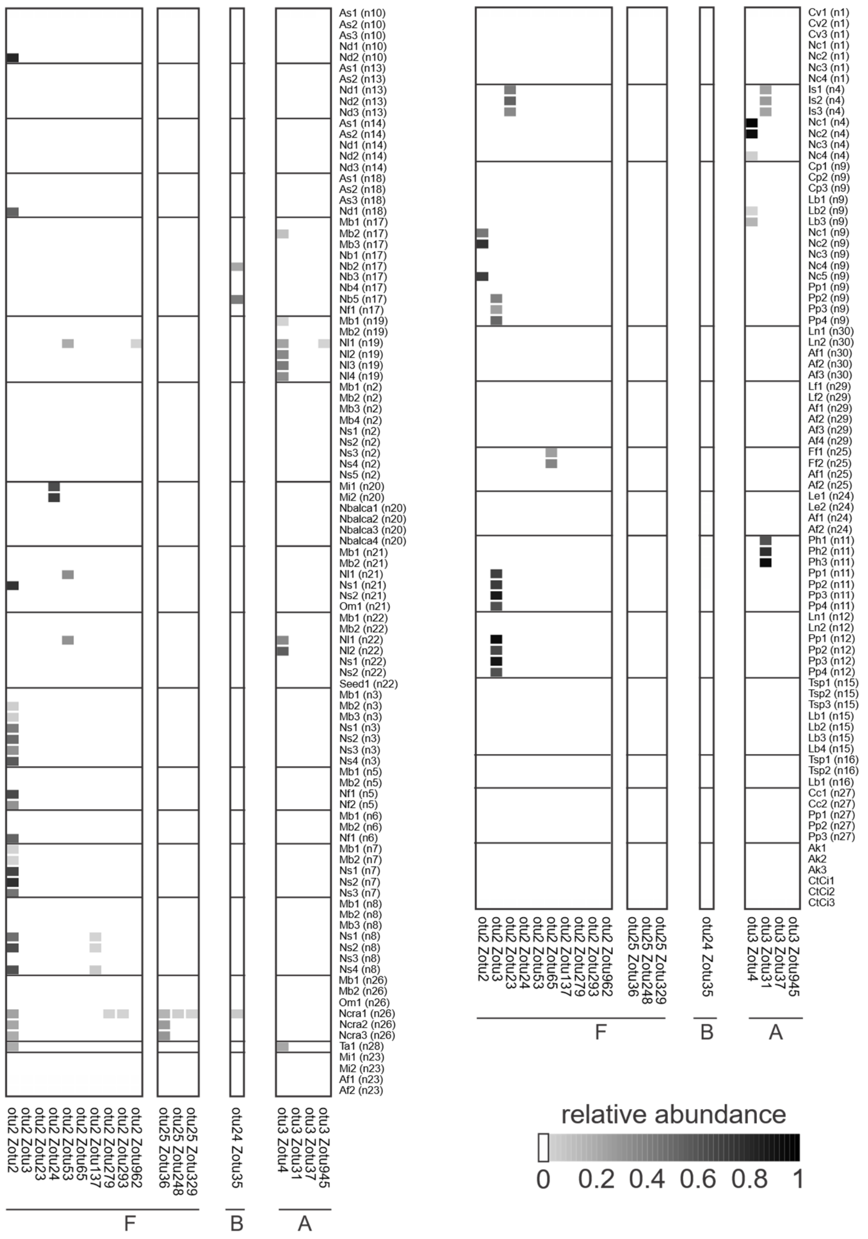


**Figure S5. Maternally transmitted *Wolbachia* bacteria are pervasive in our insect panel.** Ant and silverfish individuals are grouped according to nest origin. The relative abundances of *Wolbachia* bacteria are visualized (see bottom right). *Wolbachia* zOTU lineages are divided based on their apparent supergroup affiliation (A, B, or F). Insect specimens are indicated by their species name abbreviation and nest origin (latter between brackets).

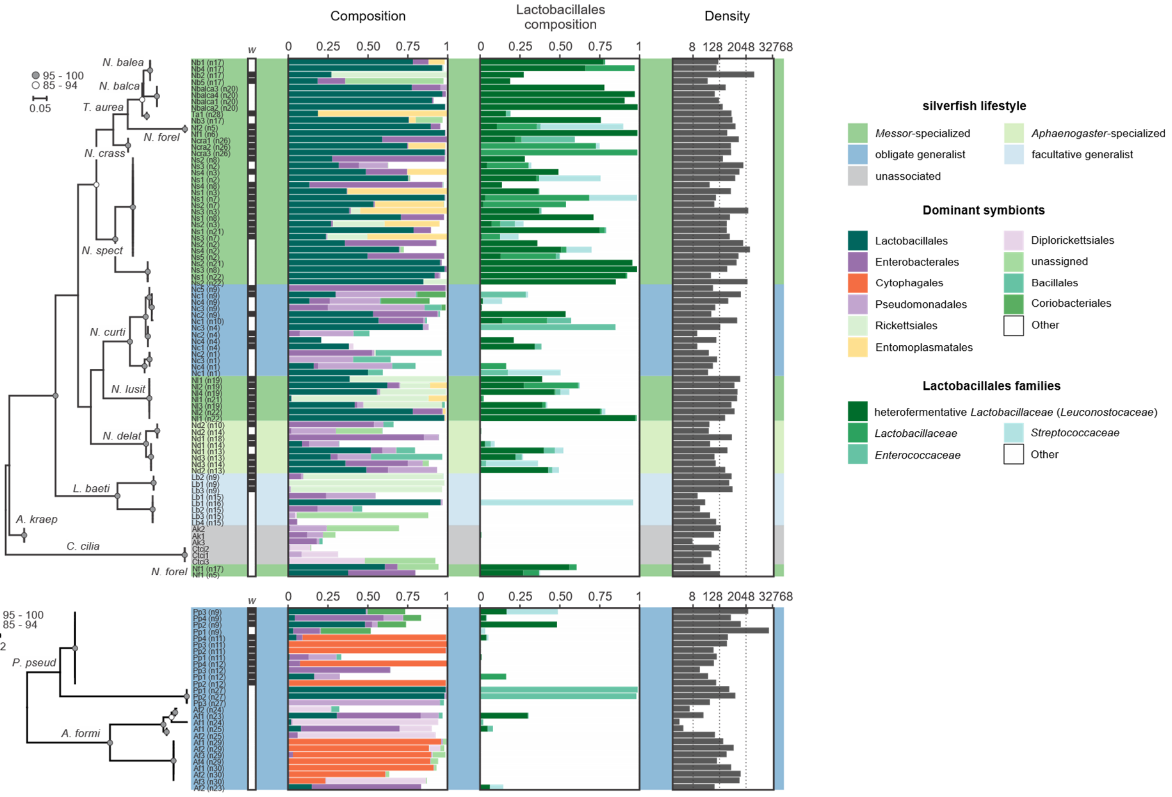


**Figure S6. Myrmecophilous lifestyle shapes the silverfish microbiome.** This supplementary figure replicates Figure 3 but adds the insect specimen ID to each microbiome. Specimens are indicated by their species name abbreviation and nest origin (latter between brackets). The Lepismatinae and Atelurinae phylogenies are shown. *COI* amplification was not successful for three silverfish samples and their symbiotic communities are depicted below their respective phylogenetic trees. Silverfish lifestyle is colour-coded (see right). *Wolbachia* infection is indicated by a black background within the row labelled with ‘*w*’. The unit for the microbiome density is the number of symbiotic rRNA copies per ng DNA. *Leuconostocaceae* is currently considered as a later synonym of *Lactobacillaceae*.

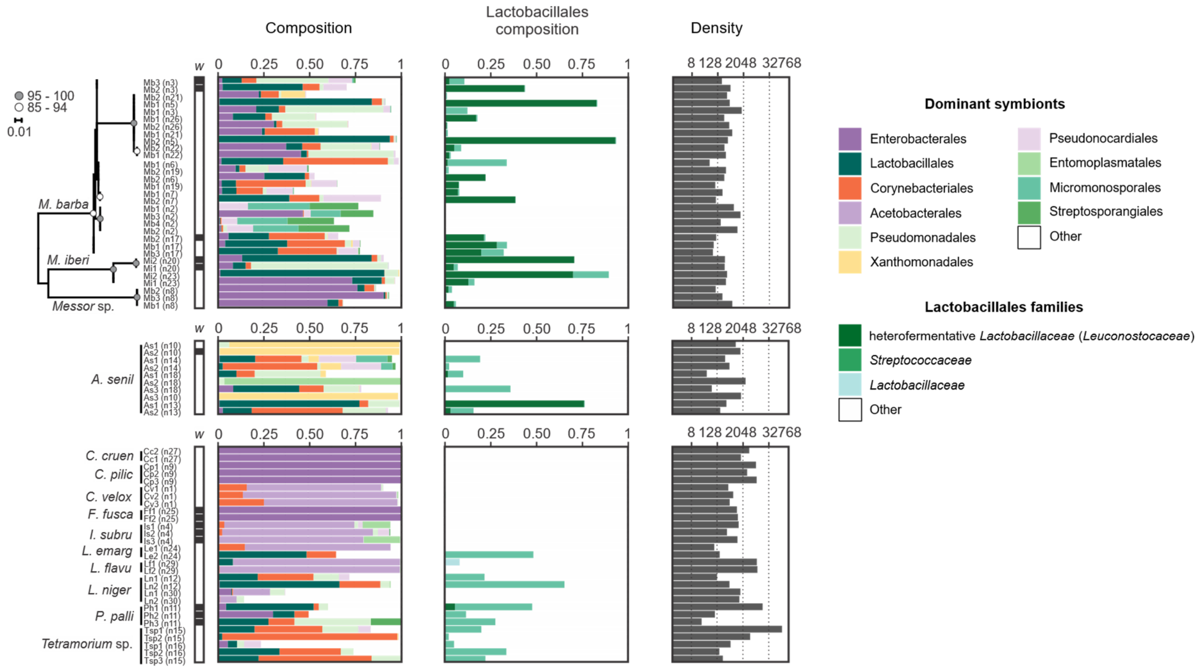


**Figure S7. Heterofermentative *Lactobacillaceae* symbionts infect *Messor* ants.** Specimens are indicated by their species name abbreviation and nest origin (latter between brackets). *Messor* ant samples are ordered according to phylogeny. *Wolbachia* infection is indicated by a black background within the row labelled with '*w*'. The unit for the microbiome density is the number of symbiotic rRNA copies per ng DNA. *Leuconostocaceae* is currently considered as a later synonym of *Lactobacillaceae*.

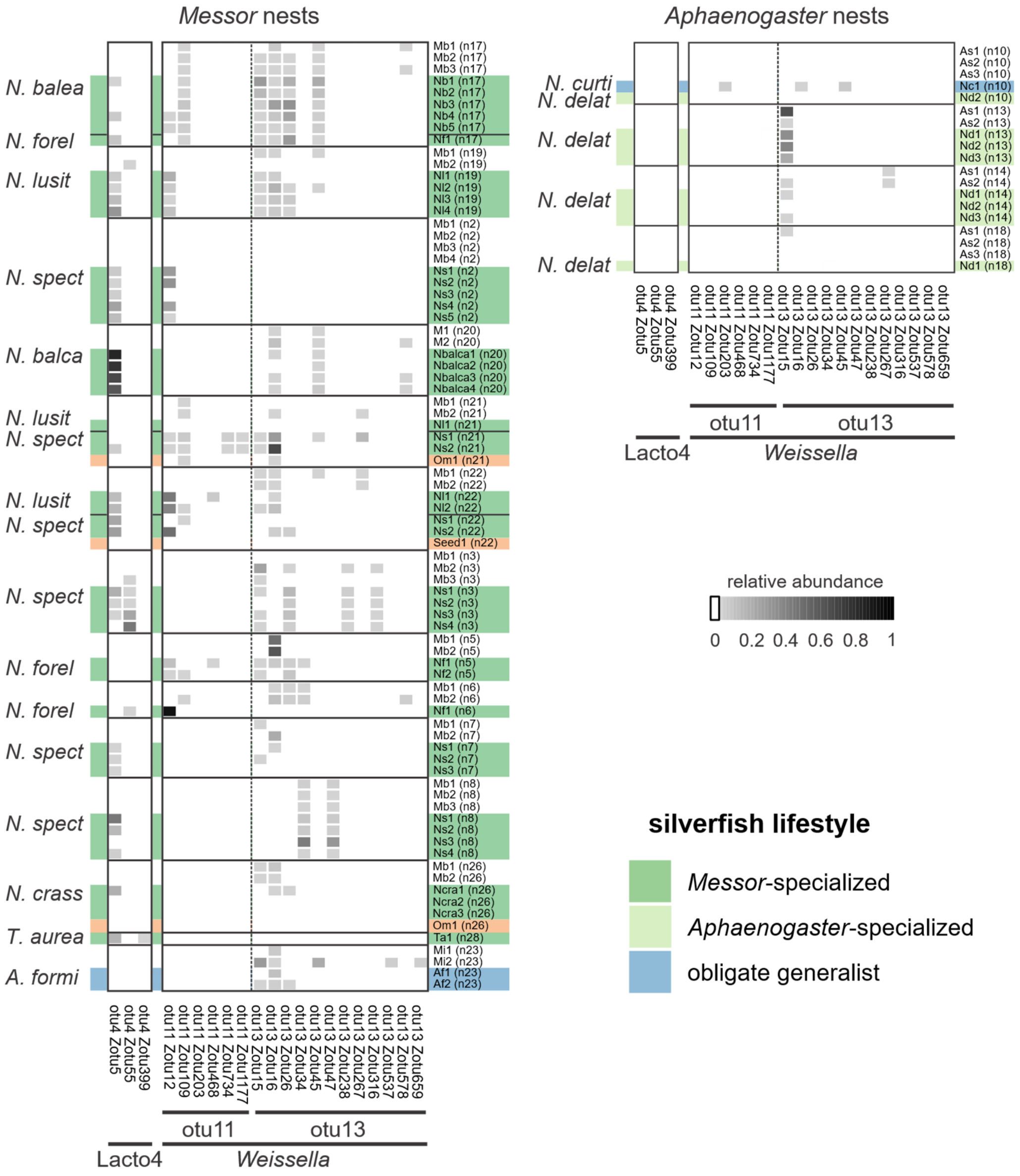


**Figure S8. Ant nest determines the infection profile of *Weissella* bacteria in *Messor*-specialized silverfish.** This supplementary figure largely duplicates Figure 4 but now also adds the insect specimen ID's. Specimens are indicated by their species name abbreviation and nest origin (latter between brackets). Ant and silverfish individuals are grouped according to nest origin. The relative abundances of heterofermentative *Lactobacillaceae* symbionts are visualized (see middle right). Silverfish lifestyle is colour-coded (bottom right). Plant seed material is indicated with an orange background. Silverfish species are further identified by their abbreviated species name. Here, OTU26 (zOTU38) was removed as only a single *C. pilicornis* colony exhibited infection (Figure S9).

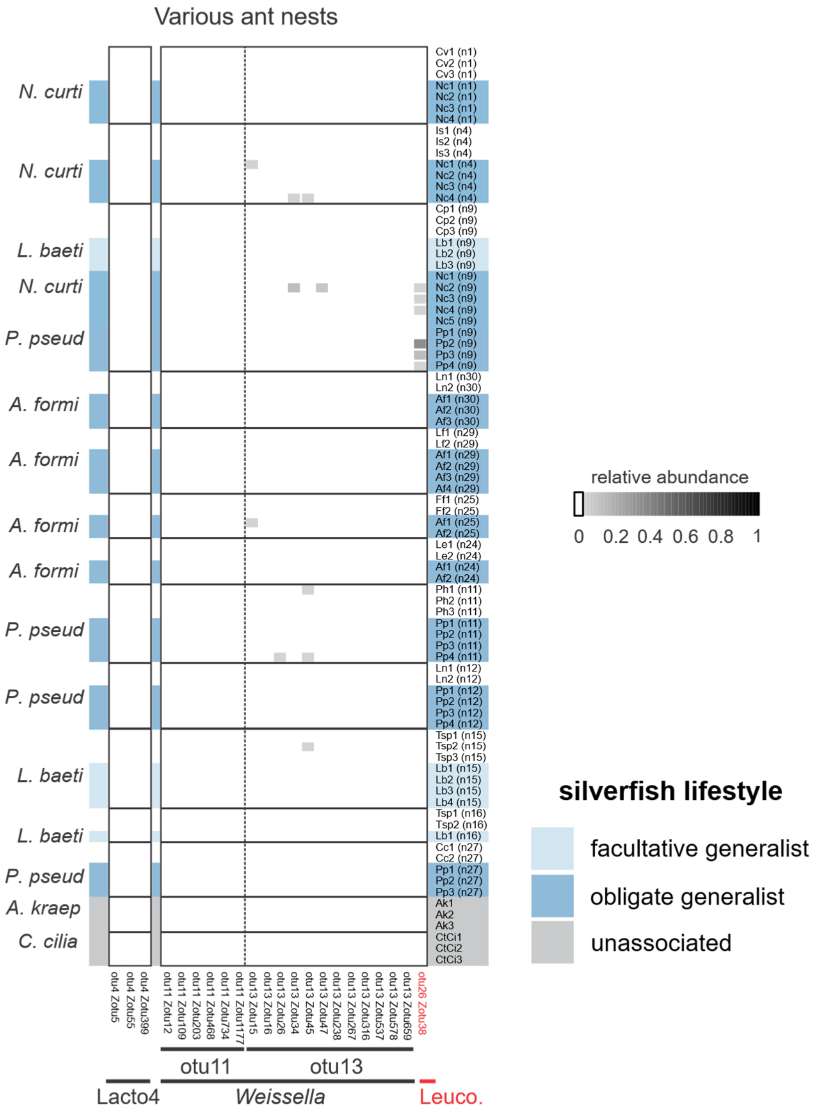


**Figure S9. Heterofermentative *Lactobacillaceae* symbionts are generally restricted to host-specialized *Neoasterolepisma* within our silverfish panel***.* Ant and silverfish individuals are grouped according to nest origin. The relative abundances of heterofermentative *Lactobacillaceae* symbionts are visualized (see middle right). Silverfish lifestyle is colour-coded (bottom right). Silverfish species are further identified by their abbreviated species name. Here, OTU26 (zOTU38) was restricted to a single *C. pilicornis* colony (depicted here in red font).
